# Strong environmental memory revealed by experimental evolution in static and fluctuating environments

**DOI:** 10.1101/2023.09.14.557739

**Authors:** Clare I. Abreu, Shaili Mathur, Dmitri A. Petrov

## Abstract

Evolution in a static environment, such as a laboratory setting with constant and uniform conditions, often proceeds via large-effect beneficial mutations that may become maladaptive in other environments. Conversely, natural settings require populations to endure environmental fluctuations. A sensible assumption is that the fitness of a lineage in a fluctuating environment is the time-average of its fitness over the sequence of static conditions it encounters. However, transitions between conditions may pose entirely new challenges, which could cause deviations from this time-average. To test this, we tracked hundreds of thousands of barcoded yeast lineages evolving in static and fluctuating conditions and subsequently isolated 900 mutants for pooled fitness assays in 15 environments. We find that fitness in fluctuating environments indeed often deviates from the expectation based on static components, leading to fitness non-additivity. Moreover, closer examination reveals that fitness in one component of a fluctuating environment is often strongly influenced by the previous component. We show that this environmental memory is especially common for mutants with high variance in fitness across tested environments, even if the components of the focal fluctuating environment are excluded from this variance. We employ a simple mathematical model and whole-genome sequencing to propose mechanisms underlying this effect, including lag time evolution and sensing mutations. Our results demonstrate that environmental fluctuations have large impacts on fitness and suggest that variance in static environments can explain these impacts.

## Introduction

The increasing scale of experimental evolution, thanks to new levels of throughput and parallelism, provides unprecedented opportunities to understand evolutionary dynamics and mechanisms. Unifying principles have emerged about evolution in a single environment, including that replicate populations display parallel adaptation in recurrent genes, operons, or pathways^1–5^, with successive mutations having a saturating effect on fitness, resulting in declining adaptability over time^6,7^. However, laboratory studies have mostly examined static environments, which have relatively temporally constant and spatially uniform conditions, and which often select for mutations that exhibit costly fitness tradeoffs if conditions change^8–12^. Such tradeoffs are relevant in natural settings where populations must endure distinct and intermittently harsh conditions, unlike in continually replenished laboratory environments.

Experimental evolution in fluctuating environments, while relatively uncharted, is making strides toward bridging this gap. One review found that costly fitness tradeoffs arise less often in heterogeneous than homogenous environments^13^. Evidence also suggests that fluctuating environments produce more surprising and unpredictable results. At the population scale, fluctuations have been shown to lead to more variation across evolving replicates^14,15^. At the individual level, a mutant’s fate can be sensitive to the timescale^16^ and order^17^ of fluctuations, and the molecular basis and fitness of mutations emerging from fluctuating environments can be different from those in static environments^18,19^. These findings align with theory that predicts that evolution is less deterministic in fluctuating environments^20,21^.

The variety of evolutionary outcomes seen in fluctuating environments raises the possibility that fluctuations render an environment fundamentally different from the sum of its parts. If fitness is additive in a fluctuating environment, the fluctuations might be decomposed into static components, and fitness in each component simply averaged to predict the overall fitness in the fluctuating environment. However, a theoretical study demonstrated a counterintuitive finding that temporal fluctuations do not always time-average fitness effects^20^. Supporting this result, there is growing empirical evidence of non-additive fitness in fluctuating environments. For example, a study of yeast gene-deletion mutants in two fluctuating environments found many instances of non-additivity, due to both to under- and over-performance in fluctuating environments^16^. Another study highlighted the fallibility of fitness assays across environmental gradients in predicting fitness in fluctuating conditions for a range of organisms, finding inconsistency between parameters obtained in static environments and performance in fluctuating environments^22^. While these results suggest that fluctuating environments may be inherently different from static environments, the mechanisms of non-additivity in fluctuating environments remain undefined.

One difference between fluctuating and static environments is the nature of the transitions they contain. For example, in laboratory batch culture, microbes grow from low density until depleting the resource, after which the culture is diluted into fresh medium, re-starting the cycle. If the medium is the same every cycle, we consider this a static environment. Yet even in such ‘static’ conditions, the dilution step requires a transition from a state of dormancy to growth, manifesting as a lag time. Batch culture evolution of yeast in a static glucose-limited environment led to the emergence of adaptive mutants that gained fitness advantage in the lag phase, with the magnitude of this advantage determined by the growth parameters of the previous cycle^23^. Their fitness advantage was thus accrued during one cycle but realized during the next. This result suggests that a mutant’s lag time can depend on its pre-conditioning environment and raises the question of how fitness would change in a fluctuating environment, where medium composition changes from one cycle to the next. We expect lag phase to change if the carbon source changes, given that lag times between carbon sources were observed in classical studies of diauxic shifts^24^, and have more recently been associated with growth tradeoffs^25–27^ and specificity of gene expression^28^. Additionally, epigenetic processes mediate phenotypic plasticity in fluctuating environments^29–31^, generating cell-to-cell variation that promotes survival in transiently stressful conditions^32,33^. Environmental fluctuations could thus alter fitness for a variety of reasons.

Here, we investigate how differences between transitions in fluctuating and static environments cause changes in fitness using high-throughput genome barcoding. We first evolved clonal populations of *Saccharomyces cerevisiae*, each containing ∼500,000 unique barcoded lineages, in static environments, including stressors and alternate carbon sources, as well as in temporally fluctuating pairwise combinations of these environments. We tracked barcoded evolutionary trajectories and subsequently selected a pool of 889 mutants for follow-up fitness assays in all environments. By comparing fitness in fluctuating environments to the average fitness in the component static environments, we see a spectrum of non-additivity. Furthermore, frequency trajectories of individual mutants reveal striking changes in fitness in the components of fluctuating environments compared to the corresponding static environments. These changes are often hidden in overall fitness, to some extent, since they partially cancel each other. We show that these changes reveal a pattern based on fitness differences across static environments, whereby mutants with larger variance in fitness are more sensitive to transitions and exhibit more environmental memory, or fitness change in fluctuating environments. We employ a simple mathematical model and whole-genome sequencing to propose mechanisms underlying this environmental sensitivity, including that mutations that cause higher fitness variance are environment-specific and may shift lag times more in fluctuating environments. Our results demonstrate that environmental fluctuations alter fitness and fundamentally differentiate dynamics in static from fluctuating environments.

#### Definitions

<u>Non-additivity</u>: Difference between overall fitness of a mutant in a fluctuating environment and its time-averaged fitness in the component static environments.

<u>Environmental memory</u>: Difference between fitness in an individual component of a fluctuating environment and fitness in a static environment with this component only.

## Results

### Large dataset acquired through experimental evolution and fitness assays

To study how evolution and fitness depend on the environment, we began by evolving barcoded yeast in static and fluctuating environments. We propagated clonal populations containing ∼500,000 uniquely barcoded lineages, as developed in a previous study^34^, for ∼168 generations in serial batch culture. We carried out evolutions in five static environments and five fluctuating pairwise combinations of these static environments (**Fig. 1A**, **Table 1**, Methods; we included the remaining five pairwise combinations in later fitness assays). Static environments contained either glucose (*Glu*), *S. cerevisiae*’s preferred carbon source, or less-optimal alternate carbon sources (*Gal*: galactose, *Lac*: lactate), and in two environments we supplemented the glucose condition with stressors known to affect *S. cerevisiae* fitness^35,36^ (*NaCl*: sodium chloride, *H_2_O_2_*: hydrogen peroxide) (**Table 1**).

**Figure 1:**
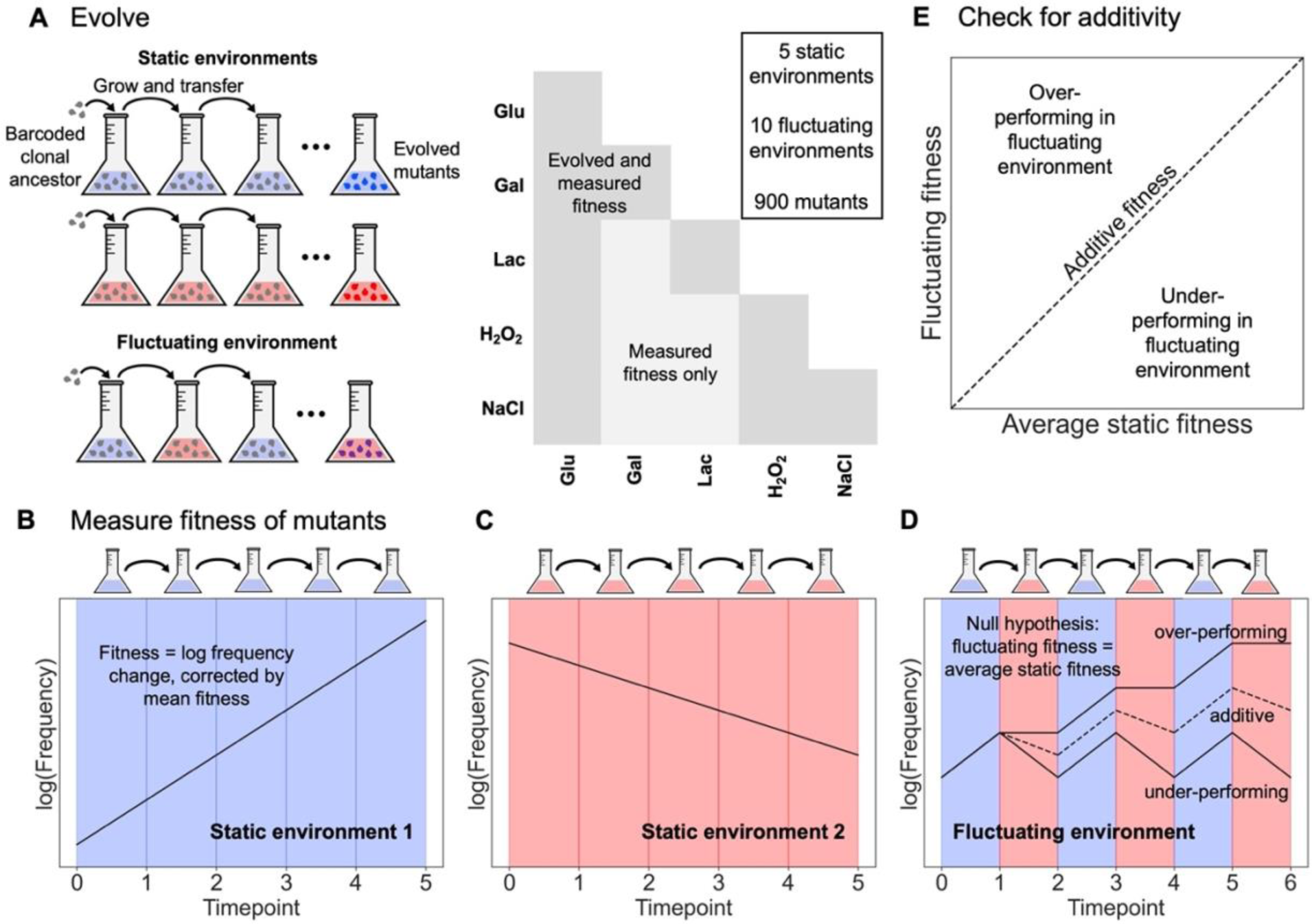
Fitness assays test for non-additivity in mutants evolved from static and fluctuating environments. **A** We evolved the same barcoded library in five static conditions and five fluctuating pairs of these static conditions. We then isolated 889 mutants from the evolution experiments and performed pooled fitness assays in all environments, plus five additional fluctuating environments. **B-C** After correcting for mean population fitness, we estimate a mutant’s fitness from the slope of its log frequency. **D** The null hypothesis of additive fitness in a fluctuating environment is represented by the average of the slopes in panels B-C. Non-additive fitness might result if transitions in the fluctuating environment alter fitness of a mutant. **E** Non-additivity can be visualized by plotting observed fitness in a fluctuating environment vs. the null hypothesis. Mutants with non-additive fitness will appear above or below the one-to-one line.

**Table 1:** Descriptions of static and fluctuating environments.

| Environment | Composition | Evolved | Fitness measured | Multi-hit genes (no. of hits) |  |
| --- | --- | --- | --- | --- | --- |
| Static: |  |  |  | Homozygous SNPs | Copy number variation |
| Glu | SC + 1.5% glucose | • | • | ZAP1 (2) |  |
| Gal | SC + 1.5% galactose | • | • | ISN1 (2) |  |
| Lac | SC + 0.7% lactate | • | • | - |  |
| H<sub>2</sub>O<sub>2</sub> | SC + 1.5% glucose + 2.5mM hydrogen peroxide | • | • | - |  |
| NaCl | SC + 1.5% glucose + 0.5M sodium chloride | • | • | tT(AGU)B (2), PAU8 (2) | ENA1 / ENA2 / ENA5 (18) |
| Fluctuating: |  |  |  |  |  |
| Glu/Gal |  | • | • | ISN1 (2) |  |
| Glu/Lac |  | • | • | ARN2 (2), GRX3 (2), ISN1 (2) |  |
| Glu/H<sub>2</sub>O<sub>2</sub> |  | • | • | AFT1 (14), BOL2 (7), SNF5 (2), ZRT3 (2), RRP5 (2), ISN1 (2) |  |
| Glu/NaCl |  | • | • | - | ENA1 / ENA2 / ENA5 (7) |
| H<sub>2</sub>O<sub>2</sub>/NaCl |  | • | • | - | ENA1 /<br>ENA2 /<br>ENA5 (4) |
| Gal/Lac |  |  | • | no evolution | no evolution |
| Gal/ H<sub>2</sub>O<sub>2</sub> |  |  | • | no evolution | no evolution |
| Gal/NaCl |  |  | • | no evolution | no evolution |
| Lac/ H<sub>2</sub>O<sub>2</sub> |  |  | • | no evolution | no evolution |
| Lac/NaCl |  |  | • | no evolution | no evolution |

We tracked changing barcode diversity during evolution and subsequently chose timepoints for reviving and isolating mutants (Fig. S1). We created a pool of 889 uniquely barcoded mutants for fitness assays in all static and fluctuating environments. We added the pool at 5% relative abundance to a preculture containing the ancestor strain, giving each barcoded lineage a starting frequency of ∼5*10^-5^. We measured lineage frequencies over several cycles (eight generations per cycle) to determine fitness relative to the ancestor (**Fig. 1B-D**; Methods). Briefly, we define fitness as log frequency change, corrected by mean population fitness:

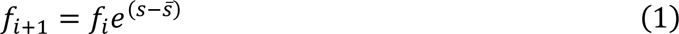

Here, *f*_*i*+1_ represents frequency of a lineage at timepoint *i* + 1, which changes exponentially from timepoint *i* if its fitness *s* differs from mean population fitness *s̅*, which we measure with barcoded neutral strains. We infer fitness in fluctuating environments separately for each component and define the overall fitness as the average of the two components.

To check the reproducibility of fitness assays, we included three biological replicates of each condition. Replicates are strongly correlated (Figs. S2-S3), but we exclude one of the ten fluctuating environments, *Lac*/*H_2_O_2_*, in analyses summarizing environments, because replicates in this environment are weakly correlated (Fig. S3, Methods), but include it individually (Figs. S3, S6-S7). To focus on mutants adaptive in their home evolution environments, we exclude those with fitness less than 0.05 in their home environment, or per-cycle fitness 5% greater than ancestral fitness, but results do not change for a higher threshold or for no threshold (Fig. S4). Unsurprisingly, most mutants are adaptive in their home environment: excluding mutants below the adaptive threshold brings the size of the pool to 695 (78% of the initial pool). Adaptive mutants evolved in fluctuating environments often have lower fitness in their home environment compared to mutants evolved in static environments (Fig. S5A). There are exceptions, such as the *Glu*/*H_2_O_2_* environment, but highly adaptive mutants often evolved in *Lac*, *H_2_O_2_*, and *NaCl*, suggesting that static environments with stressors or a non-fermentable carbon source lead to large fitness gains.

### In fluctuating environments, non-additive fitness and memory are common

To determine whether fitness in fluctuating environments is additive, we compare it to fitness in static environments, where we observe a diversity of behaviors within the pool (Fig. S5). The null hypothesis of additive fitness in a fluctuating environment follows from the expectation that mutants will increase or decrease their frequency in a given condition regardless of the previous condition from which they were diluted. If mutants instead have non-additive fitness, this may indicate that transitions between different conditions have unique effects on mutants’ growth compared to transitions between the same condition (**Fig. 1D**). Indeed, previous studies have shown that environmental fluctuations can lead to changes in fitness, either due to changes in the lag phase of growth^27,28^ or other mechanisms^16,37^ (**Fig. 1E**).

We find that non-additivity, or deviation from the time-average, is extremely common in our mutant pool (**Fig. 2A**). For brevity, we highlight three of the ten fluctuating environments, which include all five static environment components, and which contain examples of mutants both over-performing and under-performing their time-averaged fitness (see Fig. S6 for the remaining seven). To distinguish non-additivity from the effects of measurement noise, we compare observed non-additivity to that expected based on the standard error of fitness in static environments across all timepoints of biological replicates (a less conservative approach uses just the standard error across replicates; see Methods). We find that 82% of fitness measurements in fluctuating environments deviate from the time-average more than expected from measurement noise. The magnitude of non-additivity is greater than 0.25, or the median fitness of the pool in static environments, in 7% of cases, showing that fitness often deviates from additivity by a large magnitude.

**Figure 2:**
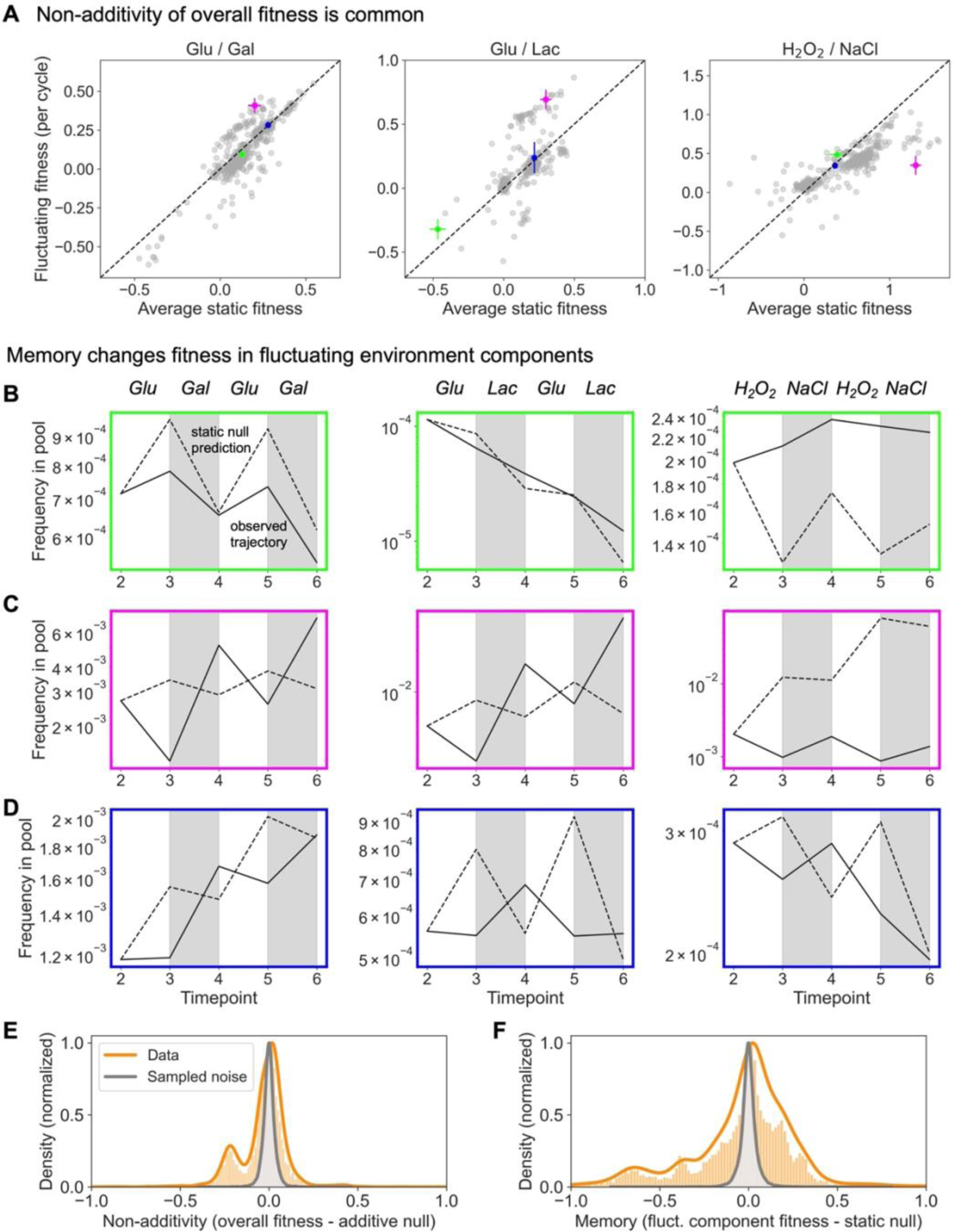
Non-additivity is common, but it masks even greater fitness changes. **A** Plotting fitness of the mutant pool in three fluctuating environments against the null hypothesis of the average of their fitness in the static environments reveals many cases of non-additivity, or differences between these two quantities. We highlight particular cases in green, magenta, and blue. **B** Frequency trajectories (solid lines; mean of three replicates) show that the static null prediction (dashed lines; also calculated from replicate mean) is inaccurate, and that overall non-additivity masks the totality of fitness change, or environmental memory. **C** Frequency trajectories of non-additive mutants show reversals of the outcome expected based on static environment fitness. **D** Frequency trajectories of additive mutants also show reversals, even if the overall trajectory and fitness is similar to what was expected. (Note the finer scale in these plots compared to those in panel **C**.) **E** Non-additivity, or the difference between overall fluctuating fitness and the additive prediction, is plotted for all measurements in nine fluctuating environments (orange). Sampling from prediction uncertainty (Methods) shows that the measurement exceeds noise. **F** Memory, or the difference between fitness in fluctuating environment components and corresponding static environments, is greater in magnitude than non-additivity.

Even more striking than non-additive behaviors, mutants show large changes in fitness within components of fluctuating environments that may not be captured by non-additivity. To fully quantify the difference between static and fluctuating environments, we define *environmental memory* as the difference between observed fitness in each component of a fluctuating environment and fitness in the corresponding static environment. We expect zero memory if the fitness is fully defined by growth during the current cycle, but not if fitness is influenced by (i.e. ‘remembers’) the previous cycle. We highlight several mutants with varying degrees of non-additivity that have dramatically different behaviors from expectations based on static environment fitness, leading to environmental memory **(Fig. 2B-D)**. Observed frequency trajectories in the fluctuating environment (solid lines), excluding the first two cycles, are plotted against null expectation based on fitness in the component static environments and mean population fitness at each timepoint (dashed lines; Methods). In these plots, non-additivity appears as deviation from the overall slope of the dashed line, while memory appears as deviation from segments of the dashed line in individual components.

For example, the green plots in panel **B** (corresponding to green dots in panel **A**) show examples of mutants that have nearly additive fitness but exhibit memory, or slight changes within each component that are not fully captured by non-additivity. Panel **C**, on the other hand, shows mutants with highly non-additive fitness. In the leftmost plot, a mutant with high fitness in glucose and low fitness in galactose dramatically reverses its fitness in the fluctuating components, decreasing in frequency during the glucose cycle and increasing in frequency during the galactose cycle. The other plots in panel **C** show similarly dramatic fitness reversals. Fitness reversals even occur for mutants that have additive fitness (panel **D**; note the finer scales in these plots). In this case, overall trajectories reach the expected frequency, hiding memory in the overall measurement, since over-performance in one half of the environment cancels under-performance in the other half. These results show that non-additivity in fluctuating environments is not only common, but that even in cases where the overall fitness across the whole fluctuating experiment is similar to the expectation, the component fitnesses often deviate sharply from the null expectation.

Unlike non-additivity, environmental memory includes the difference between observed and null in both halves of the fluctuating environment. Non-additivity is significant compared to measurement noise (**Fig. 2E**) and has smaller magnitude than memory (**Fig. 2F**), which also exceeds measurement noise and extends to roughly twice the range. As before, to estimate measurement noise, we sample from a normal distribution with a standard deviation equal to the standard error of predictions (Methods), finding that memory exceeds that expected by measurement noise in 87% of fitness measurements in fluctuating environments (compared to 82% of non-additivity). The magnitude of memory in either component of a fluctuating environment is greater than 0.25 in 29% of cases (this is true of non-additivity in 7% of cases), illustrating that transitions in fluctuating environments lead to radical and widespread changes in fitness, making fluctuating environments inherently different from a sequence of static environments.

### Fitness variance in static environments correlates with memory in fluctuating environments

Overall patterns of environmental memory reveal a tradeoff in which mutants gain fitness in one half of a fluctuating environment but lose fitness in the other half. To visualize memory of the pool in each fluctuating environment, we first plot fitness in pairs of static environments, coloring mutants by fitness difference (**Fig. 3A**; distance from the equal-fitness dashed line). As shown in **Fig. 3B-D**, darker shades of blue denote higher fitness in the first environment, and darker shades of red indicate higher fitness in the second environment. We preserved this shading when plotting memory, or difference in fitness between each component of a fluctuating environments and the corresponding static environment. As shown in **Fig. 3E**, the null hypothesis, or no memory, sits at the origin of these plots. Data not on the origin can still represent additive fitness if it sits on the *y* = −*x* line, because in this case, fitness change in one half of a fluctuating environment exactly cancels change in the other half. We observe that not only do the measurements generally deviate from the origin, but that they tend to cluster around the *y* = −*x* line, showing that fitness gains occur in one component of a fluctuating environment and losses occur in the other, and that they often match each other in magnitude (**Fig. 3F-H**).

**Figure 3:**
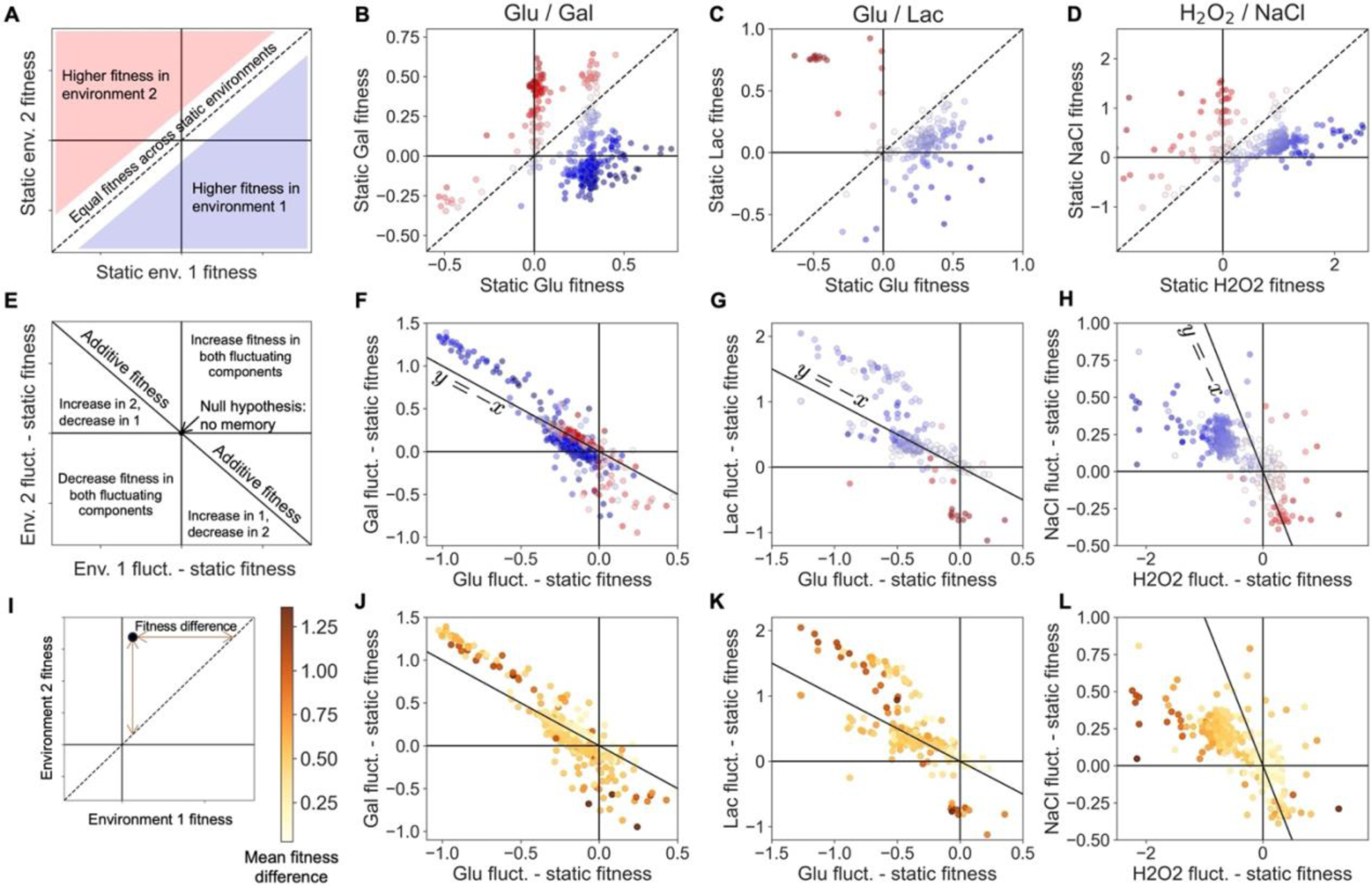
Fitness in one component of a fluctuating environment influences fitness in the other component. **A** Plotting fitness across a pair of static environments indicates mutants that have higher fitness in the first environment (blue) and mutants that have higher fitness in the second environment (red). **B-D** Fitness of the pool is shown in three pairs of static environments. **E** Plotting the difference of fitness in one component of a fluctuating environment and fitness in the corresponding static environment shows memory, or how fitness changes in fluctuating environments. The *y* = −*x* line indicates mutants that change their fitness but are still additive. **F-H** The change in fitness in the two components of fluctuating environments is negatively correlated, indicating that mutants gain fitness in one component and lose fitness in the other. The presence of blue dots in the upper left quadrants and red dots in the lower right quadrants indicates that fitness gains often occur in the environment with lower static fitness and fitness losses often occur in the environment with higher static fitness, suggesting that in a fluctuating environment, fitness is influenced by the previous condition. **I** We define fitness difference between two static environments as the distance from the equal-fitness line. **J-L** Mutants with high mean fitness difference across all ten pairs are farther from the origin in these plots, meaning that they have more environmental memory.

This negative trend appears in all ten fluctuating environments (Fig. S6), indicating that opposing fitness changes that are not fully captured by non-additivity are widespread and generic. Furthermore, mutants tend to gain fitness in the component corresponding to the static environment in which they do worse and lose fitness in the component corresponding to the static environment in which they excel, indicated by the recurrent pattern of blue dots in the upper left quadrant and red dots in the lower right quadrant. This surprising pattern suggests that memory is indeed caused by influence of the previous condition.

A possible explanation for one cycle of a fluctuating environment influencing fitness in the next is that a mutant may evolve to optimize transitions in a particular environment, whether static or fluctuating. In that case, a lineage’s physiological state at the end of one cycle might determine its transition to lag phase and growth in the following cycle, and thus lead to different consequences if the environment in the following cycle is the same or different. For mutants with bigger fitness differences across static environments, transitions between environments could foreseeably become more severe, leading to more changes in behavior in a fluctuating environment. Indeed, the color arrangement in **Fig. 3F-H** suggests that fitness difference across a pair of static environments may correlate with fitness change when those two environments fluctuate. For example, dark red and dark blue dots, representing mutants with bigger fitness differences, tend to sit farther from the origin, indicating that they have more memory. There are some exceptions to this pattern, such as the dark red and blue dots clustered near the origin in **Fig. 3F**. These are mutants with large fitness differences in static *Glu* compared to static *Gal*, and which change their fitness in *Glu/Gal* relatively less than other mutants with similar fitness differences. However, calculating global mean fitness difference, rather than just local *Glu*/*Gal* fitness difference, reveals that these mutants have low global mean fitness difference. Overall, lower global mean fitness difference translates to less memory in fluctuating environments (or placement closer to the origin in these plots) (**Fig. 3I-L**).

As a more rigorous test of whether global mean fitness difference correlates with memory in a fluctuating environment regardless of local fitness difference, we calculate partial correlations of memory in each fluctuating environment with global mean fitness difference, controlling for and excluding local fitness difference. The partial correlations are positive and significant in nine out of ten fluctuating environments (all except for *Lac*/*H_2_O_2_*), even if static environments corresponding to the components of the focal fluctuating environment are excluded from the mean fitness difference (Fig. S7, Methods). This result shows that mutants with high fitness variance are uniquely sensitive to transitions in fluctuating environments and that a particular setting is not necessary to identify this characteristic sensitivity.

The result that bigger fitness differences in static environments propagate to bigger fitness changes in fluctuating environments can be summarized by plotting mean fitness difference across all pairs of static environments against mean fitness change across all fluctuating environments (**Fig. 4A**), with memory and non-additivity plotted separately in different colors. We confirm that fitness difference is an appropriate parameter to compare to fitness change by decomposing it into two independent parameters, both of which contribute toward explaining variation in fitness change in a general additive model (Fig. S8). While both memory and non-additivity correlate with mean fitness difference, memory has a nearly one-to-one correlation, meaning that in fluctuating environments, a mutant will on average change its fitness by an amount nearly equal to its average fitness difference across static environments. While this is an average effect, it could manifest in a particular fluctuating environment as a mutant reversing its fitness, as we saw in the trajectories in **Fig. 2C-D**. Additionally, the negative trends in **Fig. 3F-H** suggest that mutants compensate for fitness increases in one component of a fluctuating environment with fitness decreases in the other component, suggesting that fitness reversals could be common.

**Figure 4:**
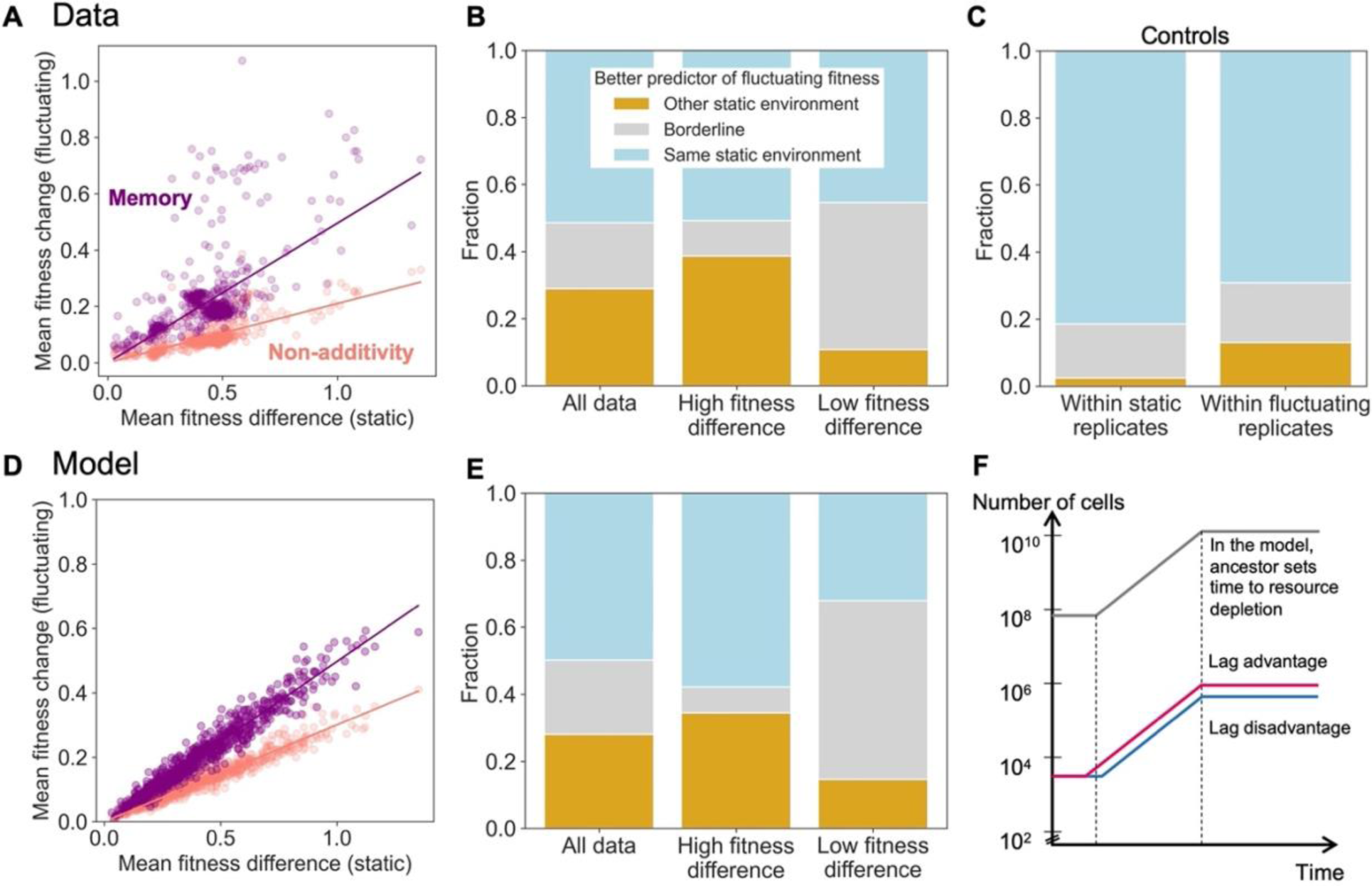
Fitness reversals in fluctuating environments are common and recapitulated by a simple model of lag time evolution. **A** Mean memory, or mean absolute difference between fitness in a static environment and fitness in the corresponding fluctuating component, correlates strongly with mean fitness difference across static environments. This nearly one-to-one correlation indicates that on average, mutants change their fitness in a fluctuating environment by an amount nearly equal to their fitness difference across static environments. Mean non-additivity, or mean difference between fluctuating fitness and average static fitness, is a lower bound of mean memory. **B** The frequency of fitness reversals in fluctuating environments is quantified by the fraction of the time that the other static environment fitness is a better predictor of fitness in half of a fluctuating environment, or 29% of overall cases. The top 10% of mutants by mean fitness difference more often reverse their fitness. **C** Fitness in one replicate of a static environment is better predicted by a replicate of another static environment than another replicate of the same static environment 2% of the time. The same is true for replicates of fluctuating environment components in 13% of cases, a noisier control because it contains half as many timepoints. **D** A simple model reproduces the correlation pattern shown in panel **A**. **E** The model reproduces the difference between mutants with high and low mean fitness difference shown in panel **B**. **F** The ancestor dominates growth in the model, where mutants’ fitness is determined by comparative lag (dis)advantages, which shift toward the mean of their distribution across static environments when the environment fluctuates.

### Mutants reverse their fitness often in fluctuating environments

To determine how often fitness reversals occur, we asked which of the two static environments is a better predictor for fitness in a component of a fluctuating environment (for example, is static *Glu* fitness or static *Gal* fitness better at predicting the *Glu* component of fitness in fluctuating *Glu*/*Gal*?). We categorize the results as the same static environment being a better predictor (e.g. static *Glu* better predicts fluctuating *Glu*), the other static environment being a better predictor (e.g. static *Gal* better predicts fluctuating *Glu*), or a borderline case if the prediction and result vary by less than 0.05 (we choose this threshold since the median standard error of fluctuating component fitness is 0.046). Overall, the other static environment is a better predictor than the same static environment in 29% of cases. In the top 10% of mutants ranked by mean fitness difference, this statistic increases to 39%, while for the bottom 10% it decreases to 11% (**Fig. 4B**). As a control, we predict fitness in static environment replicates with either other replicates of the same static environment or with replicates of another static environment; as a second control, we predict fitness in fluctuating environment component replicates with other replicates either from the same component or the other component (**Fig. 4C**). We expect both controls to be noisy, since they use replicates rather than averages of replicates, and the second control to be especially noisy, since fluctuating environment component measurements contain half as many timepoints as static environment measurements. Indeed, the second control represents the noisiest upper bound (13%) as opposed to the first control (2%). In either case, fitness reversals in fluctuating environments occur more often than would be expected due to measurement error. Most strikingly, mutants with the biggest fitness differences reverse their fitness most often, despite these reversals requiring greater change.

To explain potential mechanisms of fitness change and reversal in fluctuating environments, we employ a simple mathematical model describing our fitness assays (**Fig. 4D-F**, Methods, Supplementary Note 1). We assume that due to a starting frequency orders of magnitude higher than the mutants, the ancestor dominates growth and sets the time to resource depletion, before which mutants increase or decrease their frequencies (**Fig. 4F**). We assume that sensitivity to transitions is more likely to affect lag than growth^38^, and that mutants’ fitness changes are primarily due to changes in lag time compared to the ancestor, but that they retain the ancestor’s growth rate for a given environment. Mutants with more variance in fitness across static environments have a wider range of lag (dis)advantages. Guided by partial correlations between memory and global fitness differences as described above (Fig. S7), we also assume that in a fluctuating environment, mutants shift their lag (dis)advantages proportionately to the width of the distribution across all static environments, regardless of local fitness difference in the pair of fluctuating environments. Additionally, we eliminate differences in yield across environments as a cause of non-additivity and memory (Supplementary Note 1). These assumptions recapitulate the result that fitness change in fluctuating environments increases with mean fitness difference across static environments, and that memory exceeds non-additivity (**Fig. 4D**). We fit the overall result that the other static environment is a better predictor for fitness in a fluctuating environment component in 29% of cases by setting the shift in lag (dis)advantage to 80% of the distance toward the mean of a mutant’s distribution. The model also recapitulates that mutants with more variance in fitness reverse their fitness more often, since their larger fitness variance means the results are less likely to be borderline cases (**Fig. 4E**). These results suggest that lag time evolution could be a cause of fitness reversals in fluctuating environments.

### Environment-specific mutations may increase fitness variance and memory

In search of a genetic mechanism for significant lag time changes in mutants with high fitness variance in fluctuating environments, we whole-genome sequenced a subset of the mutant pool, obtaining 352 sequences (Methods). Most of our mutants contain only unique mutations, but several of the environments produced mutants with mutations in the same genes. Of such multi-hit genes, only five experienced more than five hits (**Table 1**). These five genes include: ENA1/ENA2/ENA5, which in tandem encode a sodium efflux pump known to be necessary for resuming growth following transfer into high salt^39^ and here displayed repeated copy number amplifications during evolution in high salt (18/43 *NaCl* mutants, 7/27 *Glu*/*NaCl* mutants, 4/34 *H_2_O_2_*/*NaCl* mutants); additionally, the genes AFT1 and BOL2, part of an iron-sensing and uptake pathway^40,41^, had SNPs in 28/72 mutants evolved in *Glu*/*H_2_O_2_* (including heterozygous SNPs). We also assayed 359 mutants for ploidy (Methods), as the DNA barcoding process sometimes induces self-diploidization^3^, categorizing 161/359 mutants as diploid. To narrow this list to pure diploids lacking other adaptive mutations, we eliminated the iron- and salt-related mutations from the identified diploids, leaving 148 “pure” diploids (note that we do not eliminate diploids with low-frequency mutations, so we cannot be sure how much of their fitness is due to these mutations vs. diploidy alone). In summary, highly repeated mutations fall into three groups: two that appear to have evolved to sense particular environments, and environment-nonspecific diploids.

We categorize these groups of mutations by plotting their mean fitness difference and mean memory. In agreement with a previous study showing that pure diploids benefit from low-variance, modest gains in fitness across a range of environments^12^, we find that diploids have moderate fitness variance and memory (**Fig. 5A**). Iron-sensing mutants have high fitness variance and memory, while salt-sensing mutants have moderate fitness variance and memory. However, when the latter group is subdivided into those with high- and low-copy number amplifications, we see that more copy number amplifications lead to comparatively higher fitness variance and memory (**Fig. 5B**), showing that more extreme forms of this environment-sensing mutation increase fitness variance and memory. These results offer a mechanistic explanation of the adaptive strategy for a subset of mutants: having an ability to sense iron or salt in their environment gives them a high fitness advantage in the relevant environment but likely comes with a costly tradeoff, such as an increased lag, when the environment changes.

**Figure 5:**
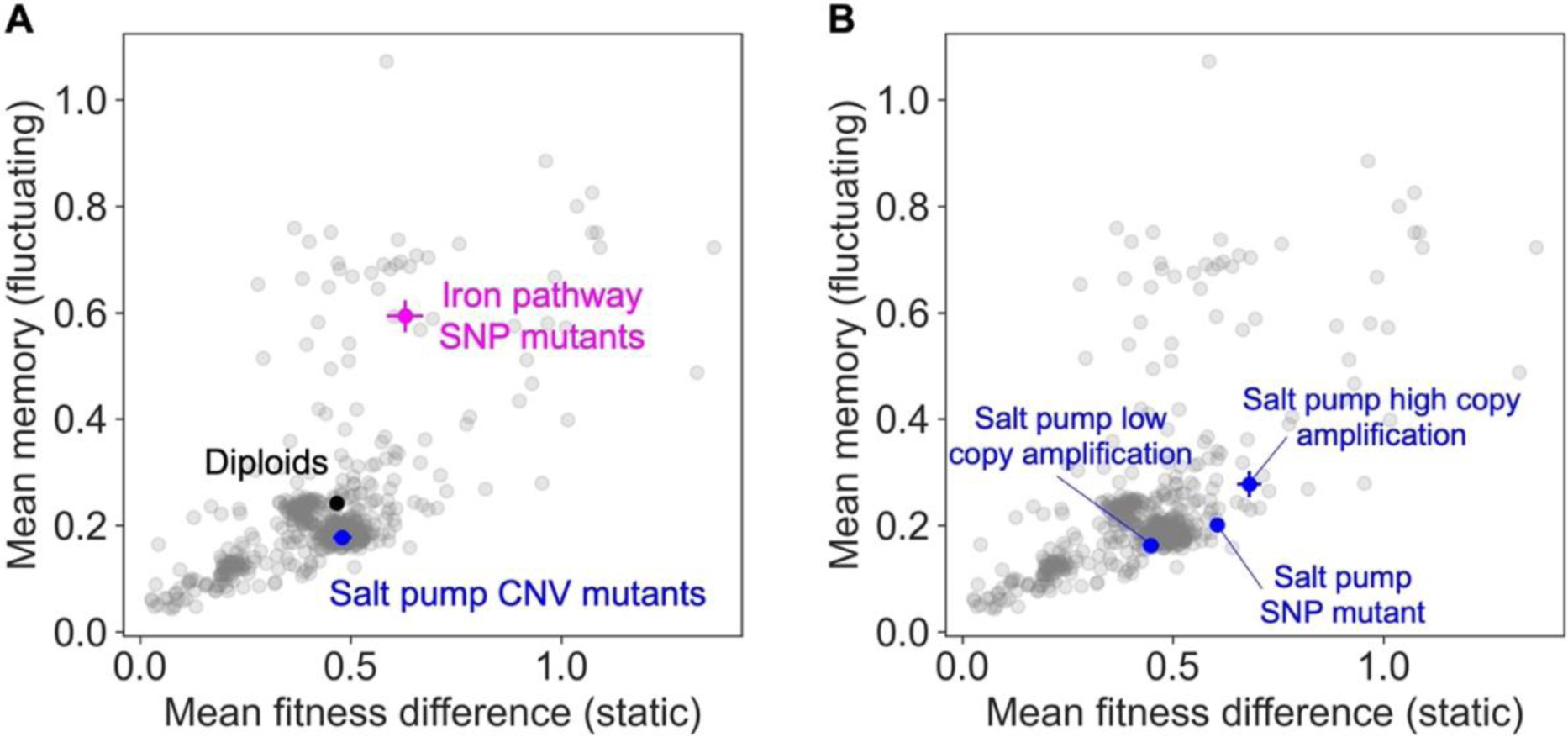
Environment-sensing mutations are associated with higher fitness variance and memory. **A** Three classes of mutants are plotted by mean fitness difference and mean memory (error bars are standard error of the mean). Diploids have moderate mean fitness variance and memory. Iron-sensing mutants have higher fitness variance and memory, while salt mutants do not. **B** Subdividing the pool of salt pump mutants into high and low copy number amplification reveals that more amplification leads to higher mean fitness variance and memory. A single SNP mutant was observed in the same set of genes as the copy number variants.

## Discussion

Natural environments are consistently perturbed by environmental fluctuations, and the amplitude and frequency of some fluctuations are expected to increase^42^. An additional concern is the ability of emerging viruses, drug-resistant bacteria, and mutating cancer cells to adapt to ever-shifting host environments. Predicting evolution in fluctuating environments is therefore important, but experiments have thus far mostly focused on adaptation to static conditions. Our study finds that fitness in a fluctuating environment can be profoundly altered by transitions that are lacking in static environments. Moreover, these changes are at least partially concealed by measurements of fitness across the complete cycle of fluctuating environments, due to a tradeoff that causes fitness to increase in one component of a fluctuating environment and to decrease in the other component. We show that mutants with high fitness variance across environments are uniquely sensitive to transitions, reversing their fitness most often in fluctuating environments, even when this variance is measured across static environments not included in the fluctuating environment components.

The observation that fitness in one component of a fluctuating environment influences fitness in the other component suggests that the physiological state of a population at the end of its growth cycle influences its growth in the following cycle. This result agrees with previous measurements of yeast in static environments, which showed that fitness accrued in the final stage of the growth cycle is realized during the lag phase of the following cycle for a set of highly adaptive nutrient-sensing pathway mutants evolved from a minimal glucose-limited medium^23^. Subsequent measurements revealed that these mutants are deleterious when the media is supplemented with 0.5M sodium chloride^12^. Based on our findings in the current study, we would expect that these glucose-sensing mutants’ large fitness variance would lead to environmental memory in fluctuations. To test this expectation, we grew those mutants in an environment fluctuating between optimal and high-salt conditions, indeed finding that their frequencies crashed in optimal conditions after exposure to high-salt conditions (Fig. S9). These results echo the environmental memory presented in this study, where we evolved and measured mutants in a complex instead of minimal medium. The agreement across differing types of media and fluctuations (both stressors and alternate carbon sources) suggests a general mechanism that causes strains, particularly those with environment-sensing mutations, to change their lag phase in one condition after exposure to another. As a result, predicting evolution in fluctuating environments based on static environments may be difficult.

While extrapolating from static to fluctuating environments may be difficult, our study also suggests that changes in fluctuating environments may be governed by a global structure whereby a strain’s fitness variance in static environments predicts its fitness change in fluctuating environments. A related phenomenon apparently detrimental to predicting evolution, epistatic interactions between mutations, has also been elucidated by similar global patterns. Originally observed in empirical studies^6,43,44^, these patterns reveal a structure whereby a strain’s background fitness predicts the fitness effect of a new mutation, leading to unifying models^45^ that have recently been applied to a novel context of predicting ecological function from species interactions^46^. The success of these global patterns in interpreting interactions between mutations and between species bodes well for the problem outlined here of interpreting interactions between environments.

To further generalize the trend that we demonstrate between fitness variance in static environments and fitness change in fluctuating environments, more investigation is needed beyond our comparison of a set of recently evolved mutants to their ancestral strain. For example, other systems such as ecological communities approaching stable states should be examined for the existence of this pattern. We grew the mutants in our pool from very low relative frequencies in the presence of a majority of ancestor cells to prevent frequency-dependent effects, but studies of frequency-dependent interactions in communities have also shown evidence of additive^47,48^ and non-additive^49,50^ effects. Our study also did not address the effects of the timescale and order of fluctuations, nor did we examine environments fluctuating between more than two conditions. In all cases, mathematical models offer a promising tool for interpreting how outcomes may be altered by fluctuation- and frequency-dependent effects^51^.

Our whole-genome sequencing results suggest that sensing mutations may lead to higher fitness variance and environmental memory. In contrast to previous studies^3,11^, however, most of our sequenced mutants contain only unique mutations. This result might reflect a diversity of adaptive mechanisms in the evolutionary conditions, the relatively short timescale of the evolution experiments, or undetected adaptations, such as aneuploidy^52^ or epigenetic modifications. Epigenetically induced phenotypic heterogeneity could contribute to a bet-hedging strategy^30,32,53^, increasing geometric fitness at the cost of arithmetic fitness across conditions^54^. If such a generalist strategy decreases fitness differences across environments, considering our results, it should also decrease memory in fluctuating environments. Our finding that mean fitness difference correlates with memory independent of local fitness difference (Fig. S7) suggests a mechanism underlying some mutants’ sensitivity to transitions and others’ lack thereof. Transcriptomic and epigenomic analysis of mutants in static and fluctuating environments could shed light on phenotypic plasticity or epigenetic mechanisms underlying memory.

Predicting the effects of increasing environmental fluctuations is an important challenge in ecology and evolutionary biology. Here we have demonstrated that the widespread existence of memory in fluctuating environments makes them qualitatively different settings than just the sum of their component static environments. This memory manifests as fitness changes in fluctuating environments, likely because the growth environment in one cycle influences lag phase in the following cycle. While transitions from one cycle to the next occur in both static and fluctuating environments, the latter impose more severe shifts, especially for mutants with greater fitness variance across static environments. This trend reveals that while null expectations about fitness in fluctuating environments are inadequate, their failure illuminates biological patterns and avenues for future investigation.

## Methods

### Long-term evolution experiments

#### Founding populations

The barcoded yeast population used in this study was constructed in a previous study^34^ and contains around half a million uniquely barcoded transformants. The library was created from an S288C derivative, BY4709, in which a ‘landing pad’ was inserted into a neutral location in the yeast genome that allows for high-frequency, site-specific genomic integration of plasmids via the Cre–loxP recombination system. We created new stocks of this library and sequenced the barcode region to confirm a high diversity of ∼500,000 barcodes before beginning batch culture experiments.

#### Evolution batch cultures

We evolved the population in ten evolution conditions (Table 1), which are composed of static or fluctuating regimes of five environments: *Glu*, *Gal*, *Lac*, *H_2_O_2_*, *NaCl*. We began evolution experiments by growing a single overnight 15-mL preculture containing the founding population in synthetic complete (SC) medium with 1.5% glucose (*Glu* media). We then transferred 400ul of this pre-culture to all evolution environments, containing 100mL of various media types. All cultures were evolved by serial batch culture in 500-ml Delong flasks (Bellco) at 30°C and 223 r.p.m. We diluted cultures every 48 hours 250X by inoculating 400ul of culture into 100 ml fresh medium. The SC base medium (per liter: 2g US Biological Drop-out Mix Complete, 6.7g RPI yeast nitrogen base) was supplemented with either 1.5% D-glucose (RPI) (*Glu*), 1.5% D-galactose (RPI) (*Gal*), or 0.7% sodium DL-lactate (Sigma) (*Lac*). In the remaining two environments, we supplemented the *Glu* medium with either 0.5M sodium chloride (RPI) (*NaCl*) or 2.5mM hydrogen peroxide (Ward’s) (*H_2_O_2_*). All media was prepared in the weeks prior to experiments and stored at 4°C until the final days before use, when we pre-warmed it in a 30°C incubator. We tested hydrogen peroxide concentration in the *H_2_O_2_* media after weeks of storage using the MQuant Peroxid-Test kit (for 1-3-10-30-100 mg/l) and found that the hydrogen peroxide had reduced to ∼60% of the original concentration. After learning of this effect, we added hydrogen peroxide directly to media during the dilution/transfer step, rather than during media preparation, in the subsequent fitness assays (described below). In contrast to short-term fitness assays, which included three biological replicates for each condition (as described below), evolution experiments contained one or two replicates. We evolved in all fluctuating environments with two replicates except for *Glu*/*Gal*, which had one replicate, and we evolved in static environments with one replicate, except for *Glu* and *NaCl*, which both had two replicates. We propagated the cultures for 20-23 growth cycles and stored 2 ml saturated culture by aliquoting 1 ml cells into 1 ml 50% glycerol in two Eppendorf tubes and storing at -80°C. The remainder of the cell culture was centrifuged and the cell pellet was resuspended in 2 ml sorbitol solution (0.9 M sorbitol, 0.1 M Tris-HCL pH 7.5, 0.1 M EDTA pH 8.0), aliquoted into two Eppendorf tubes and stored at −20°C for genomic extraction.

### Barcode amplicon and whole-genome sequencing

#### DNA extraction

We extracted genomic DNA from each sample using the Norgen Fungi/Yeast Genomic DNA Isolation 96-Well Kit, supplemented with lyticase. In collaboration with Norgen sales representatives and research scientists, we modified this kit to use 96-well plates for all steps. We extracted DNA using ∼10% of our spun-down samples, or about 10^9^ cells. Final DNA yield was assayed with a Biotek fluorescent plate reader and averaged ∼4ug per sample.

#### Barcode amplicon library preparation and sequencing

To track changes in barcoded lineage frequency over the course of evolution, we conducted PCR amplification of the barcoded region with a two-step protocol. The primers used in the first step of amplification are the same as described in ref ^12^, but with a different amplification protocol. In both steps of PCR, we performed three reactions per sample in the standard Q5 master mix (per 50ul reaction):

- 10ul 5X Q5 Reaction Buffer
- 1ul 10 mM dNTPs
- 2.5ul 10uM forward primer
- 2.5ul 10uM reverse primer
- 0.5ul Q5 High-Fidelity DNA Polymerase
- DNA
- Nuclease-free water (to 50ul)

In the first step, we aimed for 100ng of genomic DNA template (or approximately 1/1000 of total DNA in a saturated 100-ml experiment) per reaction, but since we added equal volumes to all reactions, actual amount was sometimes as low as 25ng or as high as 250ng. For the first cycle, we amplified the barcode region with the following program:

1. 98°C for 30 seconds
2. 98°C for 10 seconds
3. 62°C for 20 seconds
4. 72°C for 30 seconds
5. Repeat steps 2-4 29X for a total of 30 cycles
6. 72°C for 3 minutes
7. 4°C hold

After checking amplification for ∼25-50 randomly chosen reactions on a gel, we then diluted the step 1 product 500X into the second reaction, by diluting 25X into nuclease-free water, and then 20X into the same master mix listed above. The only difference between step 1 and 2 master mix was the primers—we used IDT for Illumina UD Indexes in the second step (https://support-docs.illumina.com/SHARE/AdapterSeq/Content/SHARE/AdapterSeq/Nextera/UDIndexesSequencesNXT.htm), with unique pairs of forward and reverse primers for each reaction to avoid error from index hopping or template switching^55^. We then ran the following program to attach the IDT indices to barcode amplicons:

1. 98°C for 30 seconds
2. 98°C for 10 seconds
3. 62°C for 20 seconds
4. 72°C for 30 seconds
5. Repeat steps 2-4 11X for a total of 12 cycles
6. 72°C for 3 minutes
7. 4°C hold

We again checked for amplification of randomly chosen reactions on a gel and then pooled step 2 product across the three reaction replicates, and then pooled samples in groups of 96 using 10-15ul per sample. We then performed a standard PCR purification protocol in one column per sample, eluting into 100ul nuclease-free water, and measured concentration with a Qubit fluorometer. Final concentration ranged from ∼100-200 ng/ul. Our samples were then sent to Admera Health (https://www.admerahealth.com/) for quality control (qPCR and either Bioanalyzer or TapeStation) and sequencing. We used 2x150 paired-end sequencing along with index sequencing reads on Illumina machines using patterned flow cells (either HiSeq X or NovaSeq X). All amplicon samples were sequenced with at least 20% genomic DNA spiked in (either whole genomes or phi-X) to ensure adequate diversity on the flow cell.

#### Whole-genome library preparation and sequencing

We prepared libraries with the Illumina DNA Prep kit (ILMN DNA LP (M) Tagmentation (96 Samples, IPB)). We followed the manufacturer’s protocol, with the exception that we used 1/5 the normal volume of reagents, allowing us to use the kit for five times the normal amount of samples, based on lab members’ and others’ success with this modification to the protocol. We sent our libraries to Admera Health (https://www.admerahealth.com/) for quality control (qPCR and either Bioanalyzer or TapeStation) and sequencing. We used 2x150 paired-end sequencing along with index sequencing reads on NovaSeq X Illumina machines.

### Tracking lineage dynamics during evolution

#### Processing amplicon sequencing data

Using BarcodeCounter2^56^, sequencing reads from evolution timepoints were first aligned to the reference barcode region using Bowtie2. Based on this alignment, both condition and lineage barcodes were extracted. Reads were split into different files based on their multiplexing tags, which were unique for each sample. We then clustered reads using Bartender^57^ with Hamming distance 2 and seed length 8. Examples of evolution trajectories can be seen in Fig. S1 and full barcode count data for all evolution experiments is available on Dryad (link TBD). To isolate mutants, we chose a single timepoint from each long-term evolution experiment where adaptive trajectories were sufficiently diverse, or where the most adaptive lineages reached a frequency of ∼10^-3^, for thawing and cell-sorting. The logic here is that we wanted to isolate a diversity of adaptive mutants, and thus did not want to choose an early timepoint where beneficial mutations had not yet occurred, or a late timepoint where one or several adaptive mutants had taken over the population.

### Isolation, selection, and pooling of mutants for fitness assays

#### Cell-sorting of thawed stocks

After choosing timepoints from each long-term evolution experiment, yeast clones were isolated from frozen glycerol samples from the chosen timepoints. Frozen stock (∼50ul, containing ∼5 × 10^6^ cells) was diluted into 500 μl PBS plus 1ul propidium iodide and used for flow cytometry sorting. Single cells were sorted into 96-well plates with 100ul YPD medium in each well. Five plates of cells (480 cells) were sorted for each evolutionary replicate (80 plates in total). Sorted cells were grown at 30°C for three days without shaking, and reached saturation by day three. Saturated cell culture (∼5ul) was removed from each well and inoculated into a different 96-well plate, with 100ul fresh YPD medium in each well. These replicated plates were grown at 30°C for two days without shaking to reach saturation, and used for subsequent barcode identification. The remainder of the saturated cell culture was mixed with 100ul 50% glycerol and stored at −80°C.

#### DNA barcode identification by Metagrid

High-throughput barcode sequencing of the plates of isolated and grown cells was performed as in (Li et al), with the exception that cell lysis was performed by boiling 20ul of saturated cell culture in 20mM NaOH. After sequencing, some wells were eliminated due to low coverage, and of those remaining, we eliminated those with less than 60% of reads mapping to the top barcode and those where the number of reads of the top barcode were equal to less than 1.5 times the number of reads of the next barcode.

#### Selection and pooling of mutants

From the ∼3700 wells in which we had identified single barcodes per well, we selected 889 clones for our mutant pool. We chose mutants with high fitness in their evolutionary trajectories based on Fitmut2^58^ inferences and ensured that no overlap existed between barcode sequences. We selected mutants meeting any of the following criteria: (i) their inferred fitness from FitMut2 was greater than 0.51 (ii) the probability that they were adaptive was greater than 0.5, (iii) they reached a final frequency of 10^-5^ (iv) they reached a frequency of 10^-5^ at the 6 cycles before the final one (to account for environments where a small number of barcoded lineages took over the population, reaching frequencies close to 1). Additionally, we added 10 randomly selected weakly adaptive mutants (defined to have probability adaptive between 0.125 and 0.5). On average, 89 clones were selected from each evolution condition, but counts ranged from a minimum of 40 (*H_2_O_2_*) to 120 (*Glu*, *Glu*/*H_2_O_2_* and *Glu*/*NaCl*) clones per condition. We revived cultures from frozen glycerol replicate plates and grew for two days in fresh SC medium before pipetting ∼10ul of each selected barcode into subpools, which we then consolidated into a final pool, weighting volumes to aim for equal frequency of all barcodes.

### Short-term fitness assays

All fitness assays were performed using three biological replicates. As described previously^3,12,23^, growth competitions were set up between a pool of barcoded mutants and a reference strain, although certain protocol details were changed. We grew three separate overnight pre-cultures in 25 mL SC + 1.5% glucose for each round of assays: 1) ancestor reference strain, 2) mutant barcode pool, 3) barcoded neutral strains with fitness equal to the ancestor. We then mixed these cultures at 93:5:2 ratio (correcting for measured OD). This ratio allows for mutants to compete against the ancestor rather than competing against each other, helps to minimize the change in average fitness throughout the competition experiment, and reduces the effect of any frequency-dependent fitness effects as well as any fitness-affecting interactions among the strains that may occur. The change in the frequency of each barcode over time reflects the fitness of the adaptive mutant possessing that barcode, relative to the rest of the population, which is primarily composed of the ancestor (we excluded later timepoints in some conditions where the ancestor proportion decreased to less than 50%; see Fig. S10). We inoculated 400ul of the mixed culture (∼5*10^7^ cells) into 100 mL of fresh media in 500 mL DeLong flasks. As in the long-term evolution experiments, this culture was then grown at 30°C in an incubator shaking at 223 RPM for 48 hr. After 48 hr of growth, 400ul of saturated culture was transferred into fresh media of the same type. In static environments, we transferred the serial dilution four more times, yielding five cycles over which to measure the rate at which a barcode’s frequency changed, and in fluctuating environments, we transferred five more times for six cycles.

After each transfer of 400ul, the left-over culture was frozen for sequencing barcodes later. To prepare this culture for freezing, it was transferred to 50 mL conicals, spun down at 3000 rpm for 5 min, resuspended in 2 mL of sorbitol freezing solution (0.9 M sorbitol, 0.1 M Tris-HCL pH 7.5, 0.1 M EDTA pH 8.0), aliquoted into two wells of a 2-mL 96-well plate, and stored at -20°C.

We performed DNA extraction and library preparation with the same protocol described above for evolution experiments, with one exception: we used restriction digest to cut the ApaLI restriction site in the middle of the reference strain’s barcode region. This step prevents the ancestor from being sequenced, affording greater sequencing coverage to the mutant pool. We performed the digest after DNA extraction and again after the second step of PCR, adding 1-2ul ApaLI (NEB #R0507L) and 5ul 10X Cutsmart to 43-ul samples and incubating at 37°C for at least 2 hr (or up to overnight).

### Fitness inferences and data analysis

#### Processing amplicon sequencing data

As with the sequencing data from evolution experiments, we first extracted barcodes with BarcodeCounter2^56^. We then used BarcodeCounter2 to map extracted barcodes to our known list of mutants in our pool.

#### Fitness inference

We inferred fitness of mutants by solving Eqn. 1 for all mutants at each timepoint for each condition and replicate: *f*_*i*+1_ = *f*_*i*_*e*^(*s*−*s̅*)^, where *f*_*i*+1_ represents frequency of a lineage at timepoint *i* + 1, which changes exponentially from timepoint *i* if its fitness *s* differs from mean population fitness *s̅*.

We measure mean population fitness at each timepoint using ∼50 uniquely barcoded neutral strains, which were isolated and identified as neutral, or having fitness equal to the ancestor, in ref TBD. Since we set the ancestor’s fitness to zero, we can solve for mean fitness by plugging the neutral strains’ frequencies *n* into Eqn. 1:

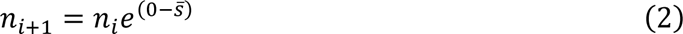

Solving for mean population fitness *s̅* gives:

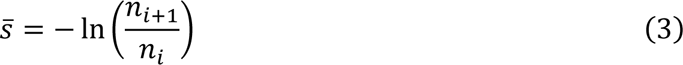

We take the average of Eqn. 3 over all neutral lineages at each timepoint *i* + 1 of each condition and replicate, and plug the result into Eqn. 1 to determine fitness of each mutant:

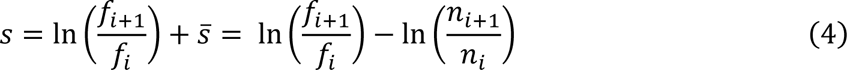

We then average this fitness across all timepoints for a given mutant/replicate/condition. We exclude the first timepoint, corresponding to the initial mixture of ancestor, neutrals, and mutants described above, from all estimates. Static environment fitness assays include five subsequent timepoints, while fluctuating environment fitness assays include six subsequent timepoints, in order grow the population in three cycles of each condition. In fluctuating environments, we exclude the second timepoint in addition to the first, assuming the first two cycles are a sort of ‘pre-culture,’ with the exception of two fluctuating environments (*Glu/H_2_O_2_* and *Lac/H_2_O_2_*) in which the barcoded pool reached a frequency greater than 50% in the final timepoint (Fig. S10). In those conditions, we excluded the final timepoint and included the second timepoint. Additionally, we excluded the final timepoint from the static environment *H_2_O_2_*, for the same reason.

In static environment inferences, we average fitness as calculated by Eqn. 4 across all timepoints, while in fluctuating environment inferences, we infer fitness separately for each of the two components and define the overall fitness as the average of the two components. Each component consists of two different timepoints, and we average these two estimates. In both static and fluctuating environment inferences, we then average fitness across all relevant timepoints in all three replicates to get a mean fitness estimate for each mutant.

We calculate standard deviation of each mutant’s fitness by assuming normally distributed errors and minimizing the negative log likelihood of a Gaussian distribution using the minimize function from the scipy.optimize module in Python, with the L-BFGS-B (Limited-memory Broyden-Fletcher-Goldfarb-Shanno with Box constraints) optimization method, setting initial guess for mean fitness to 0 and standard deviation to 0.1. To calculate standard error of each mutant’s fitness, we bootstrapped this minimization with 250 trials, sampling with replacement from all fitness measurements across all timepoints.

In rare cases, some mutants were not detected at all timepoints, either because they went extinct or just because no counts were sampled. We exclude such timepoints from our fitness inferences. In the case of static environments, if a mutant reached a detectable frequency at a later timepoint, we include this timepoint, dividing Eqn. 4 by the number of cycles between detectable frequencies. This is not possible in fluctuating environments because inferring fitness separately for the two fluctuating environment components requires measuring frequency change between consecutive timepoints only.

#### Calculating static null predictions in fluctuating environments

In **Fig. 2B-D**, we plot observed trajectories in fluctuating environments compared to predicted trajectories based on static environment fitness and mean population fitness at each timepoint (dashed lines, panels **B**-**D**). To plot these predicted trajectories, we used Eqn. 1: *f*_*i*+1_ = *f*_*i*_*e*^(*s*−*s̅*)^. Given a mutant’s known initial frequency at timepoint 2 (*f*_2_), we calculated and plotted frequency at timepoint 3 (*f*_3_) by plugging in that mutant’s fitness (*s*) in the static environment corresponding to the fluctuating environment component between timepoints 2 and 3, as well as mean population fitness (*s̅*) at timepoint 3. Mean population fitness is calculated using Eqn. 2, averaged over all neutral lineages.

#### Defining adaptive fitness

To focus our analyses on adaptive mutants, we chose an adaptive threshold of 0.05 and excluded mutants with fitness below this threshold in their home evolution environment. This threshold was based on the flattening of the curve in Fig. S4A. However, results are qualitatively the same if no threshold is used, or if an adaptive threshold of 0.1 is used (Fig. S4B).

#### Excluding data based on replicate-to-replicate correlations

We compared replicate-to-replicate correlations of fitness to identify highly variable mutants or conditions (Figs. S2-3). Correlations are strong, with Pearson r averaging 0.92 for static environments and 0.93 for fluctuating environments. We exclude the *Lac*/*H_2_O_2_* environment from this average, because fitness is weakly correlated in this environment—during experiments, growth in *H_2_O_2_* after transfer from *Lac* was not visible for >24 hours, suggesting long lag times or mortality. We exclude the *Lac*/*H_2_O_2_* environment from analyses summarizing environments, but include it individually (Figs. S3, S7-S9). Some mutants have highly variable fitness across replicates, such as the extreme differences seen across *NaCl* replicates (Fig. S2). We exclude 12 mutants with mean replicate-to-replicate fitness difference greater than 0.2 from analyses summarizing environments in **Fig. 2** and **Fig. 4**, but results do not change if they are included (Fig. S11).

#### Comparison of measurement noise to non-additivity and memory

To distinguish non-additivity and memory from the effects of measurement noise, we compared observed non-additivity and memory to that expected based on measurement uncertainty alone, as shown in **Fig. 2E-F**. To calculate the percentage of measurements of non-additivity (or memory) with magnitude greater than would be expected from measurement noise alone, we looped over all measurements of non-additivity (or memory), comparing the observed fitness to a sample from a normal distribution with mean equal to the fitness based on static environments and standard deviation equal to the standard error of this static environment fitness. The larger the error, the more likely that sampling this distribution will produce larger values of non-additive fitness (or memory) due to error alone. We repeated this loop 500 times and found the average percentage of cases in which the magnitude of non-additivity (or memory) is greater than the noise sample magnitude. In the more conservative version of this comparison (discussed in the main text), we used the bootstrapped standard error of fitness across all timepoints within all replicates, whereas a less conservative version of this comparison uses the standard error across replicates. The latter comparison has lower uncertainty (88% of cases of non-additivity have magnitude greater than the noise-induced sample magnitude vs. 82% in the former comparison; 92% of cases of memory have magnitude greater than the noise-induced sample magnitude vs. 87% in the former comparison), because the standard error across replicates averages over an entire frequency trajectory rather than including noise within each trajectory.

#### Partial correlation of fitness differences and fitness change

We calculated partial correlations of memory in a given fluctuating environment with mean fitness difference, controlling for local fitness difference in the relevant pair of static environments, using the partial_corr method in the pingouin library in Python. We first calculated mean fitness difference across nine out of ten pairs of static environments, excluding the relevant pair. For example, in the case of memory in the *Glu*/*Gal* fluctuating environment, we excluded fitness difference between static *Glu* and static *Gal*. The partial correlation is positive and significant in nine out of ten fluctuating environments (Fig. S7A; the exception is *Lac*/*H_2_O_2_*, where biological replicates are very weakly correlated).

We then excluded more pairs to see if memory in a given fluctuating environment is correlated with fitness difference across unrelated pairs of environments, controlling for fitness difference across the local pair of static environments. For example, we calculated the partial correlation of *Glu*/*Gal* memory with mean fitness difference across pairs composed of *Lac*, *H_2_O_2_*, and *NaCl*, controlling for *Glu*/*Gal* fitness difference. The correlations remain positive and significant in this case (Fig S7B), showing that we can expect more memory in unseen environments based on greater variance in fitness measured elsewhere.

#### General additive model to compare independent components of fitness difference

We used the R library mgcv to construct and fit a general additive model (GAM) to determine the effects of two independent components of mean fitness difference on mean memory. As shown in Fig. S8, fitness difference, as plotted in a two-dimensional space of fitness in two static environments, is composed of distance from the origin and angle from the equal-fitness line. We confirmed that both variables significantly contribute to variation in mean memory (origin distance estimate: 0.55, p value < 2e-16; angle estimate: 0.34, p value = 2.4e-8), and we confirmed low correlation of these variables using the check_collinearity function.

### Mathematical model of fitness assays

**Fig. 4D-E** shows the results of a mathematical model of our fitness assays. In this model, which is derived in Supplementary Note 1, we assume that the ancestor dominates the population by having a starting frequency many orders of magnitude higher than the mutants, and so sets the time to resource depletion, before which mutants can increase in frequency. Mutants have positive fitness if they grow faster than the ancestor or during the ancestor’s lag time (Eqn. 9, Supp. Note 1):

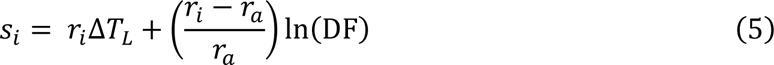

Here, *s*_*i*_ is the fitness of mutant *i*, *r*_*i*_ its exponential growth rate, Δ*T*_L_ the difference between mutant’s lag time and the ancestor’s lag time (if this quantity is positive, the mutant has a lag advantage over the ancestor), *r*_*a*_ the ancestor’s exponential growth rate, and DF is the dilution factor at each timepoint (or 250 in our experiments).

We further assume that mutants retain the ancestor’s exponential growth rate in a given environment, and fitness is determined only by lag (dis)advantage:

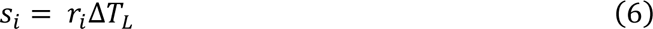

Mutants with high fitness variance have bigger differences in fitness across environments and so have wider distributions of lag (dis)advantage across environments.

Memory in a fluctuating environment component, or the difference between fitness in fluctuating and static conditions, is determined by the difference in lag (dis)advantage between conditions:

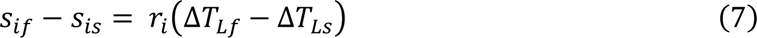

The mutant will have environmental memory if its lag (dis)advantage shifts in the fluctuating environment. Since the ancestor’s lag time likely changes in fluctuating conditions, memory will happen if the mutant’s lag time changes differently than the ancestor’s in fluctuating conditions. For example, if the ancestor’s lag is four hours in the static *Glu* and six hours in the *Glu* component of *Glu*/*Gal*, while the mutant’s lag is three hours and five hours, respectively, there is no memory. But if the mutant’s lag is three hours in both conditions, memory is nonzero and positive, meaning the mutant overperforms in fluctuating *Glu* compared to static *Glu* fitness.

To simulate fitness assays in static and fluctuating environments, we first sampled ancestor growth rates across five static environments from a random uniform distribution [0.3, 0.5]. We then sampled lag advantages for a pool of 700 mutants by first sampling each mutant’s “home” (or evolution condition origin) environment lag advantage from a random uniform distribution [0.5, 3] and setting the mean and standard deviation of its lag (dis)advantage distribution in all five static environments equal to 0.5 times its home advantage. This choice was based on the fact that we observe that mutants with higher mean fitness difference also have higher than average home environment fitness (Fig. S5E-F).

We then simulated fitness in fluctuating environments by shifting a mutant’s lag (dis)advantage away from its static environment value and toward the mean of the mutant’s distribution. Guided by our finding that memory correlates with global fitness difference when controlling for local fitness difference (Fig. S7), we shift proportionately to the width of the mutant’s distribution across all static environments. We fit the result that the wrong static environment is a better predictor for fitness in fluctuating environment components in 39% of cases by setting this shift to 80% of the distance from the local lag (dis)advantage to the mean of the distribution.

### Whole-genome sequencing analysis

We selected 359 mutants for sequencing, including as many mutants with high mean memory as possible. Two samples were excluded because their barcoded regions did not match the appropriate barcode sequence, possibly due to contamination, and an additional five were discarded due to low (<20X) coverage, leaving 352 whole-genome sequences.

#### FASTQ processing

For each sample, we received two fastq files, one for each read of the paired-end sequencing (fastqR1 and fastqR2). Reads were mapped using bwa to *Saccharomyces cerevisiae* S288C reference genome R64-1-1 (https://downloads.yeastgenome.org/sequence/S288C_reference/genome_releases/) and converted to BAM format and sorted using samtools. Read groups were fixed using the Picard tool AddOrReplaceReadGroups. Duplicates were marked using the Picard tool MarkDuplicates and the BAM file indexed using samtools.

#### Single-nucleotide polymorphism and small indel variant calling

Single-nucleotide polymorphism (SNP) and small indel variants were called using the HaplotypeCaller tool from GATK. GVCF files were then merged using GenomicsDBImport and genotyping was performed on the combined gVCF data with GenotypeGVCFs to produce a single VCF file with the called variants for all samples. Average coverage for each sample was calculated using samtools, keeping only those with at least 20X coverage. Variants were then filtered to exclude ancestral genotypes, mitochondrial variants, and using GATK default filters for other filtering. Results were checked visually in IGV. We discarded variants that appeared erroneous due to poor alignment or coverage, particularly in repetitive regions, and marked true variants as homozygous or heterozygous, based on whether the variant appeared in closer to 50% or 100% of reads. Finally, variants were annotated using snpEff.

#### Copy number variant calling

BAM files were searched for high-coverage regions with samtools and bedtools, selecting genome sites with more than 3X average coverage. To eliminate ancestral copy number variation, all selected sites were compared and results appearing in more than 15 samples were dropped. The remaining results were checked visually in IGV for copy number variation.

### Ploidy assay

We conducted a ploidy test as in ref ^3^. Saturated cultures from the 96-well plates also used for whole-genome sequencing were mixed 5ul was pipetted onto YPD + 20 ug/ml benomyl (in DMSO) on rectangular agar plates, grown at room temperature for several days and then imaged. Under these conditions, diploid growth was strongly inhibited by benomyl but haploid growth was less affected. We pipetted onto two agar plates per 96-well plate, using every other well, in order to avoid effects of nearest neighbors on colony growth.

## Acknowledgements

We are grateful to Grant Kinsler for patiently training us in the experimental system, as well as for valuable advice about PCR and sequencing, data analysis, and whole-genome sequencing analysis. We thank members of the Petrov Lab for helpful comments and suggestions. We thank Anastasia Lyulina for assistance with preliminary experiments, Sasha Khristich for guidance on molecular biology troubleshooting, Katherine Xue, Katie Solari, Katja Schwartz, and Jason Tarkington for advice on library preparation protocols, and Aditya Mahadevan for advice on whole-genome sequencing analysis. For additional helpful discussions, we thank Yuping Li, Benjamin Good, Daniel Wong, Jonas Cremer, Jeff Gore, Martina Dal Bello, Maitreya Dunham, and Rike Stelkens. Research reported in this publication was supported by the National Institute of General Medical Sciences of the National Institutes of Health under Award Number F32GM145148.

