## Supplementary Material for "Strong environmental memory revealed by experimental evolution in static and fluctuating environments"

Abreu et. al., 2023

### Supplementary Figures

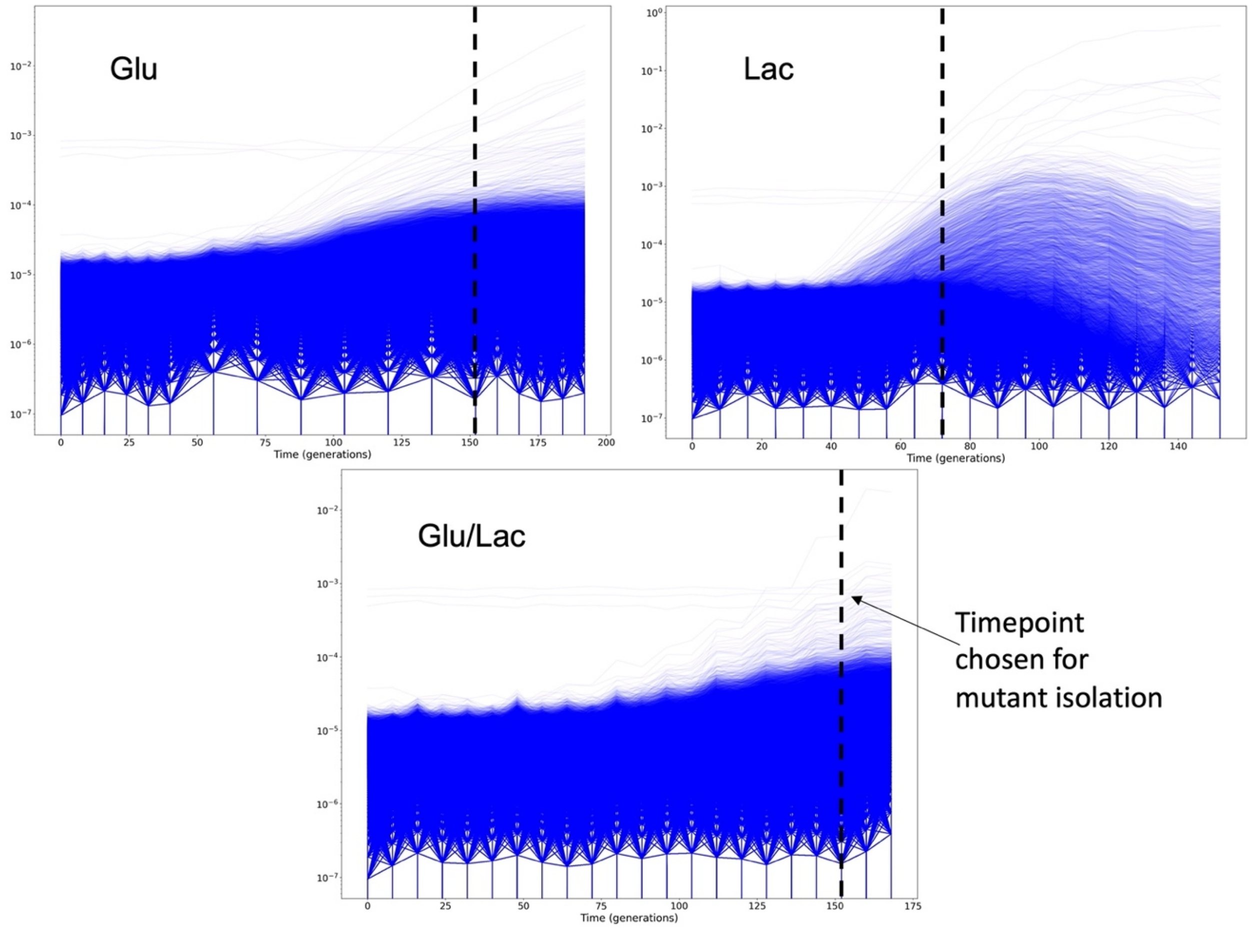


**Figure S1: Barcode trajectories show differing speeds of evolution in different environments.** Evolutionary trajectories are shown for three example experiments. We chose timepoints for isolating mutants where the most abundant lineages reached a frequency slightly above $10^{-3}$, in order to isolate a high diversity of adaptive mutants.

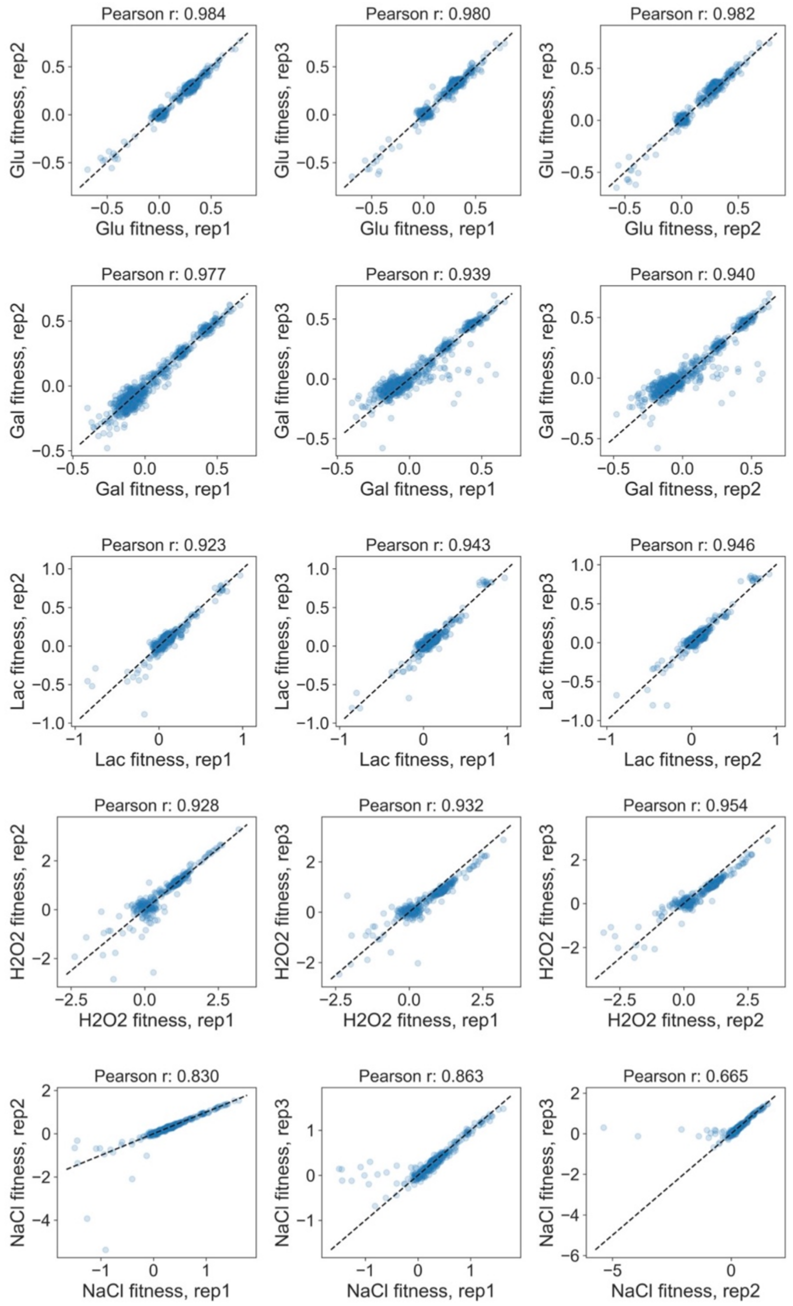


**Figure S2: Correlations between biological replicates of fitness measurements in static environments.** In static environments, Pearson correlation averages 0.92.

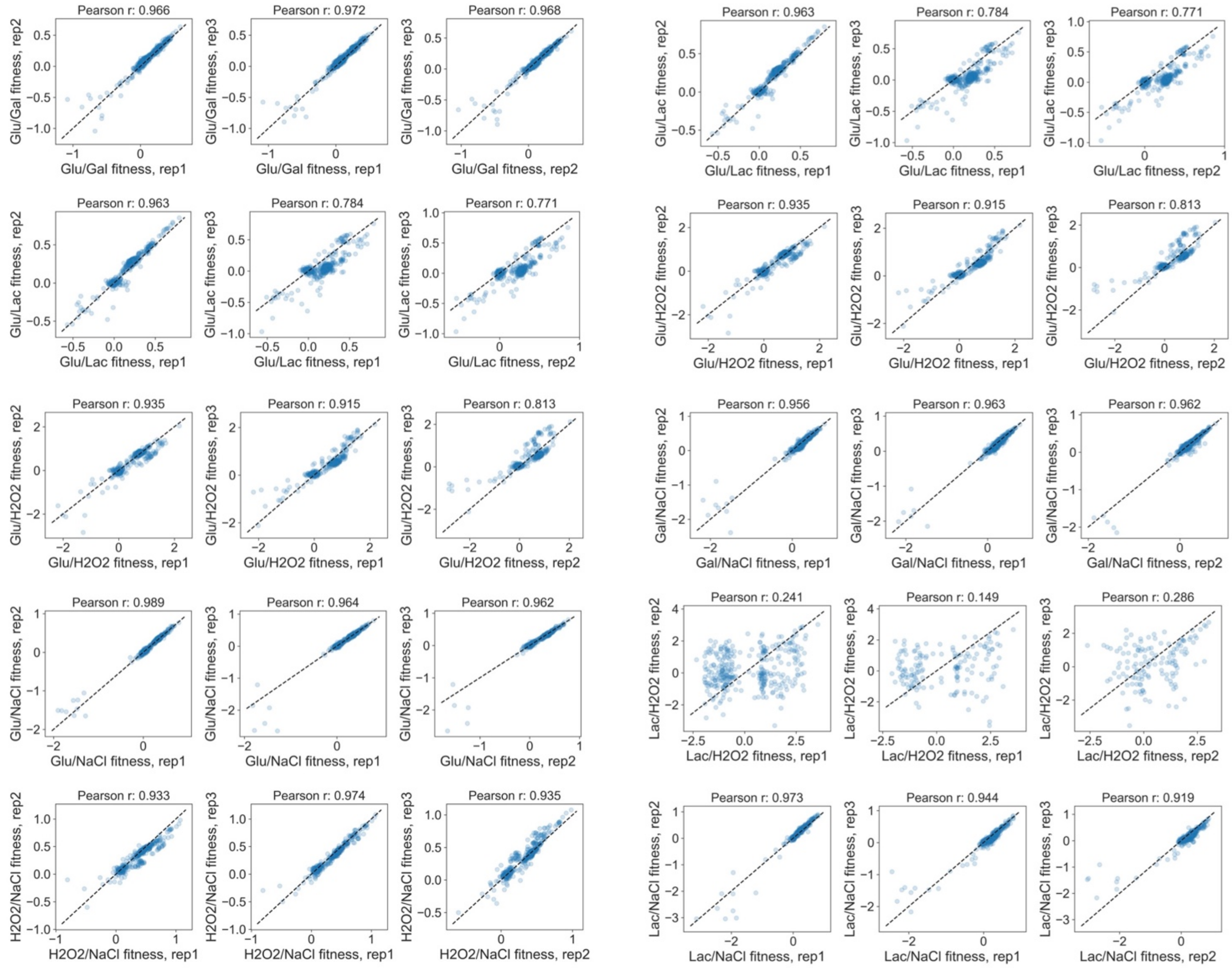


**Figure S3: Correlations between biological replicates of fitness measurements in fluctuating environments.** In fluctuating environments, Pearson correlation averages 0.93.

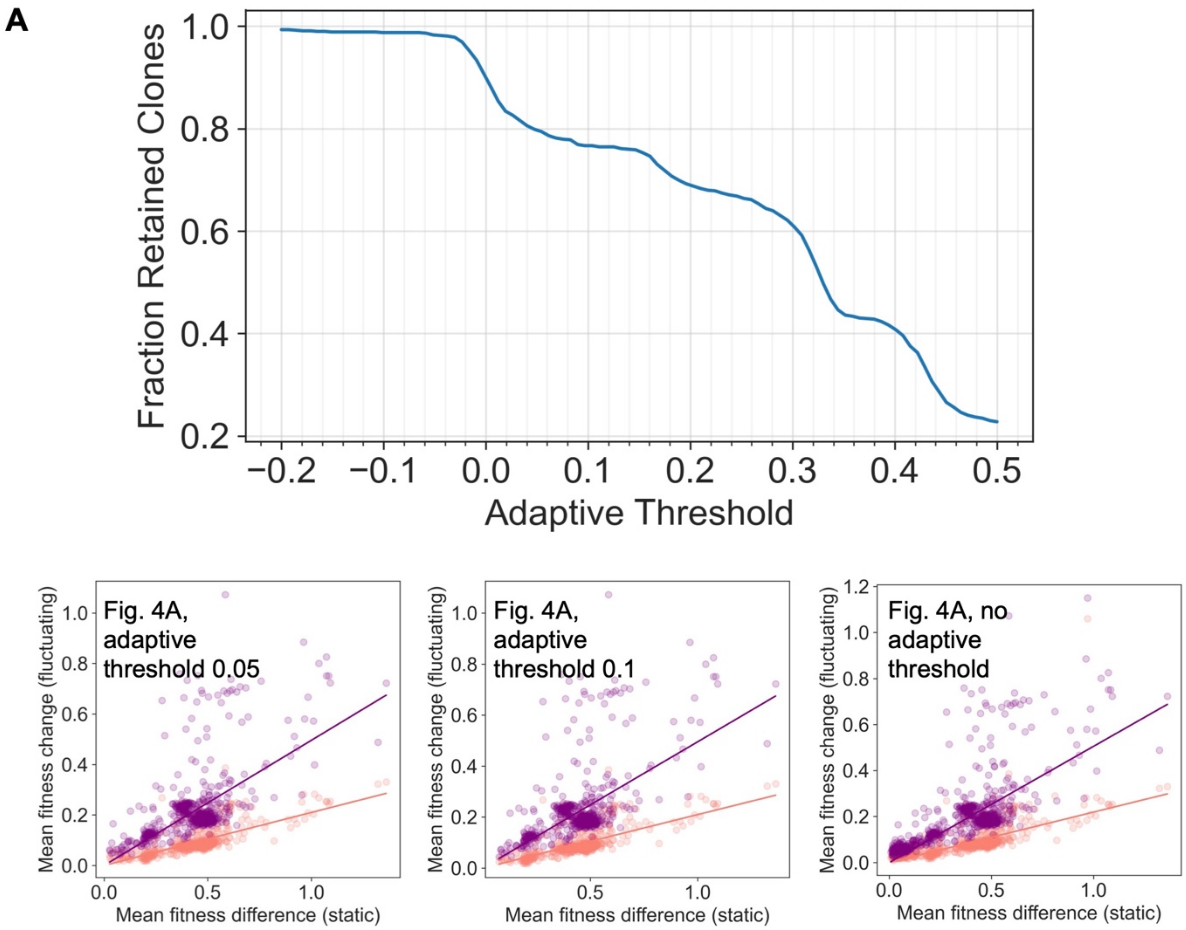


**Figure S4: Results are not sensitive to the adaptive fitness threshold used to exclude neutral mutants.** **A** We chose an adaptive threshold of 0.05 in a mutant's home evolution environment to exclude potentially neutral mutants from our analysis. **B** Using an adaptive threshold of 0.1 or no adaptive threshold produces the same qualitative final result (from Fig. 4A).

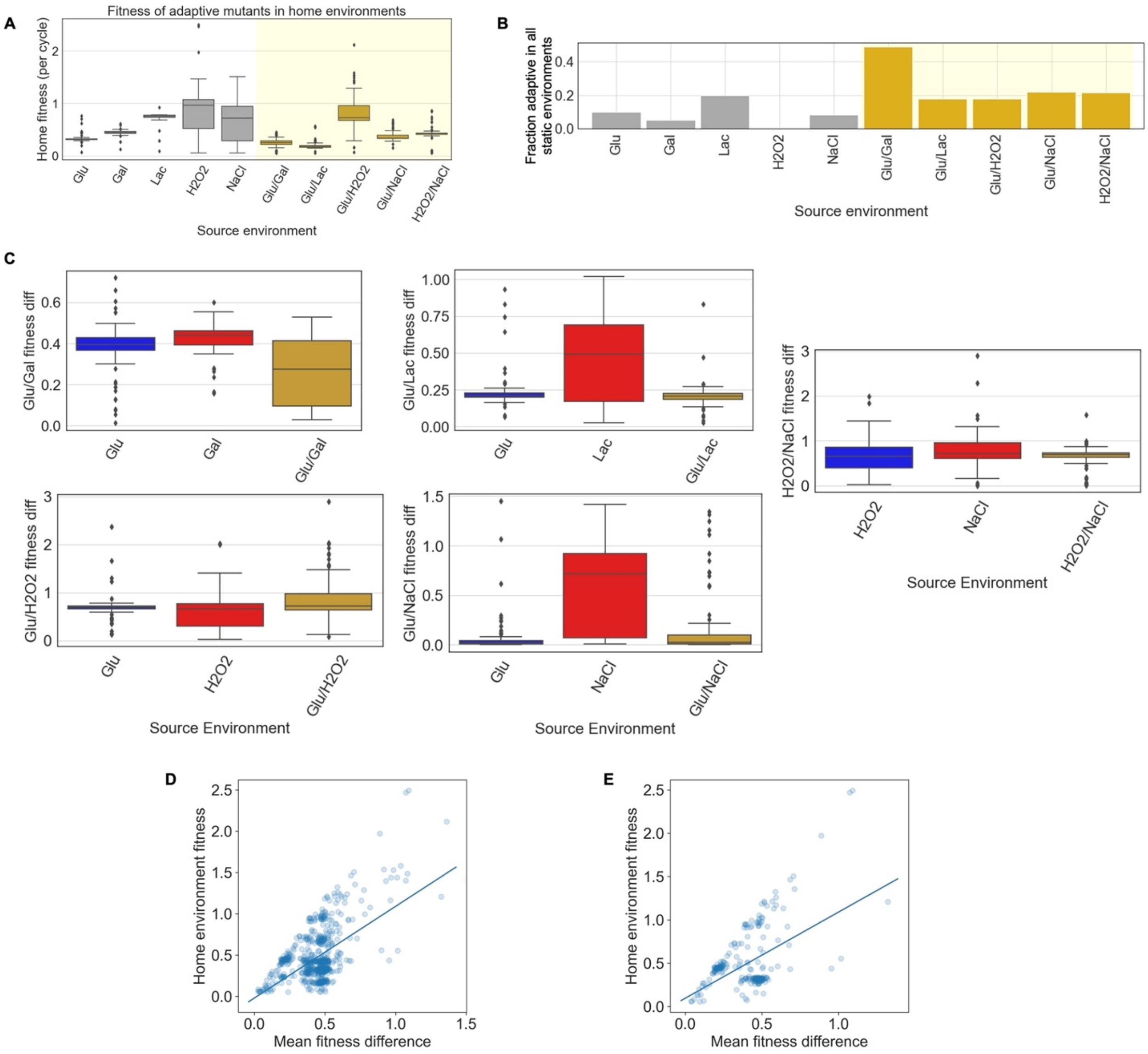


**Figure S5: Mutants evolved in static and fluctuating environments have different properties.** **A** The distribution of adaptive fitness effects shows that mutants evolved in fluctuating environments tend to have lower adaptive fitness than mutants evolved in static environments, particularly static environments with stressors or a non-fermentable carbon source. **B** Generalists, or mutants with adaptive fitness in all static environments, more often evolve in fluctuating environments. **C** Mutants evolved in a particular fluctuating environment often have lower fitness differences across the corresponding two static environments than mutants that evolved in either of the two static environments. **D** Mutants with higher adaptive fitness in their home environment tend to have a greater mean fitness difference across all static environments. **E** Mutants evolved in static environments only are shown.

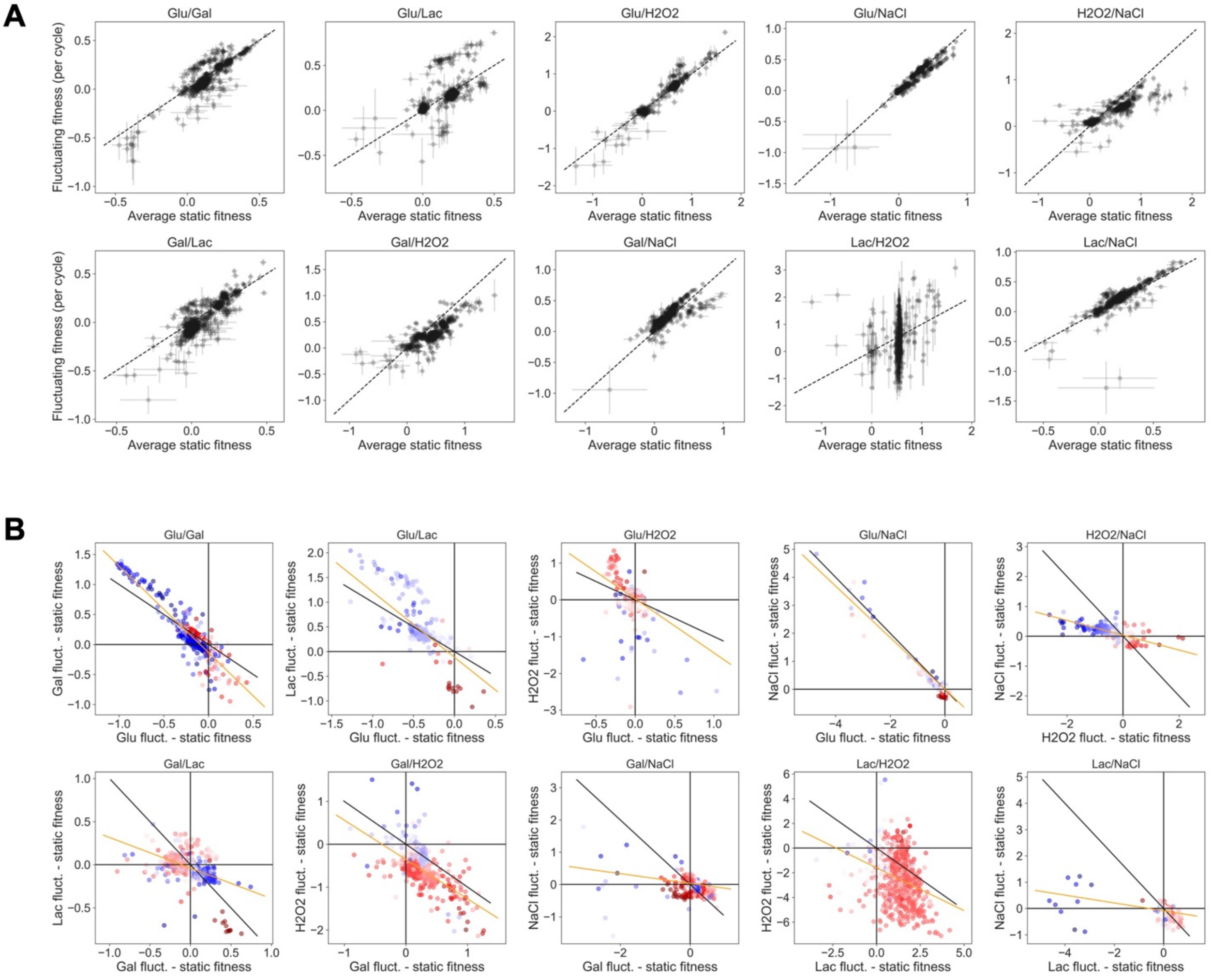


**Figure S6: Non-additivity occurs in all ten fluctuating environments, and environmental memory shows negative trend in all ten fluctuating environments. A** Non-additivity appears in fluctuating environments as data that deviates from the one-to-one (dashed) line. Error bars are calculated across all timepoints in three biological replicates by bootstrapping the maximum likelihood function (Methods). **B** Memory in fluctuating appears in fluctuating environments as a difference between fitness in a fluctuating environment component and fitness in the corresponding static environment. In the absence of memory, data sits at the origin in these plots. Data away from the origin but on the $y = -x$ line (plotted black lines) represents additive fitness, because a fitness gain in one component of a fluctuating environment exactly cancels a fitness loss in the other component. Data in all ten environments is negatively correlated, as shown by a linear regression (plotted orange lines). Data points are colored by their fitness difference across the two static environments, where blue means higher fitness in the first static environment and red means higher fitness in the second static environment.

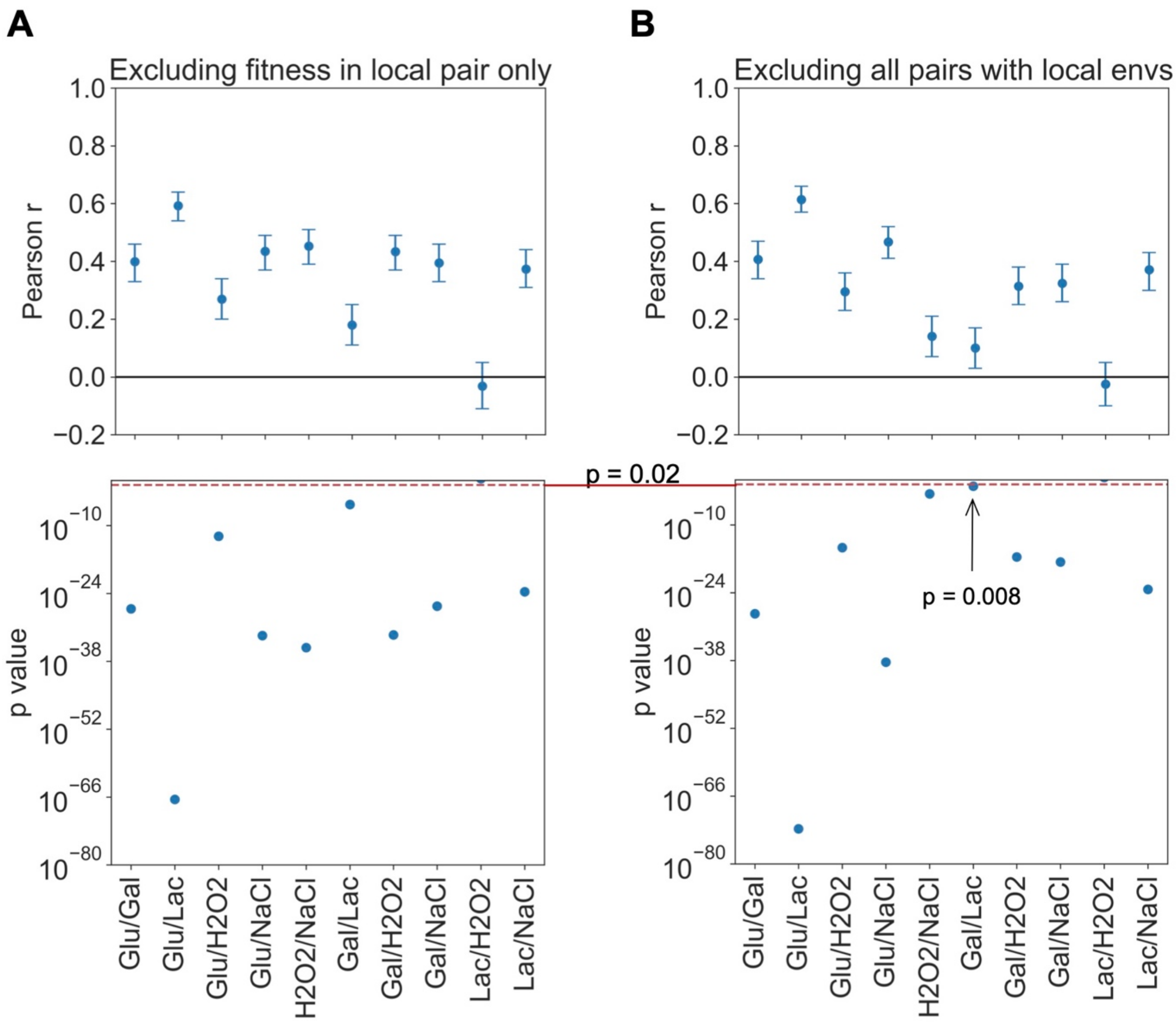


**Figure S7: Partial correlations of memory with fitness difference show that high fitness variance need not be defined locally.** **A** Partial correlations of environmental memory in each fluctuating environment with mean fitness difference across all pairs of static environments, controlling for and excluding the local fitness difference. The correlation is positive and significant in nine out of ten fluctuating environments. **B** The partial correlations remain positive and significant when excluding all pairs containing either of the static environments corresponding to the focal fluctuating components from the mean fitness.

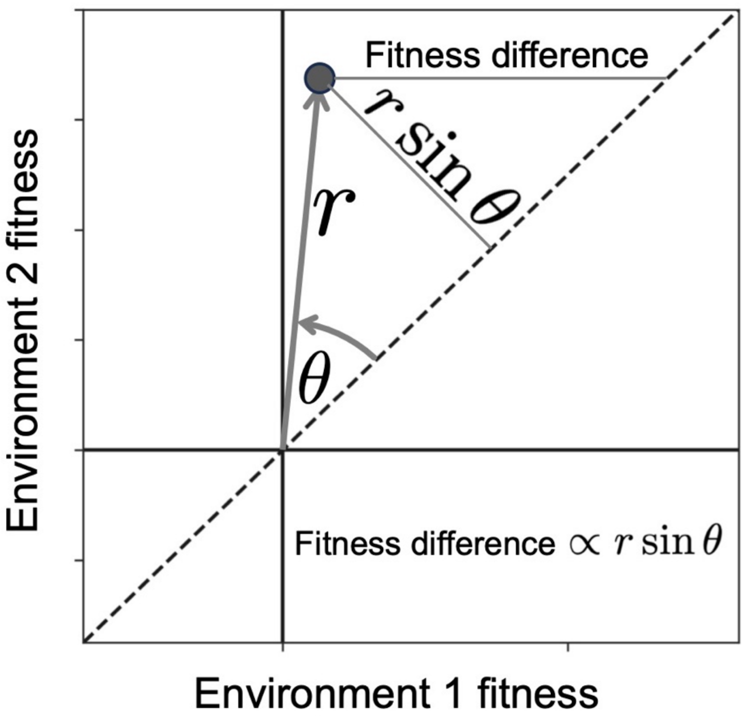


**Figure S8: Two independent components of fitness contribute toward explaining variation in mean fitness change.** Fitness difference across a pair of environments is proportional to $r * sin(\theta)$, which means that correlating mean fitness difference with a quantity such as mean memory is equivalent to correlating $r * sin(\theta)$ with the same quantity. We therefore checked whether both independent components of this quantity, $r$ and $sin(\theta)$, independently contribute toward explaining variation in mean memory, using a general additive model (GAM). We find that both contribute ($r$ estimate 0.55, p < 2e-16; $sin(\theta)$ estimate 0.34, p = 2.4e-8), justifying our use of the product of the two.

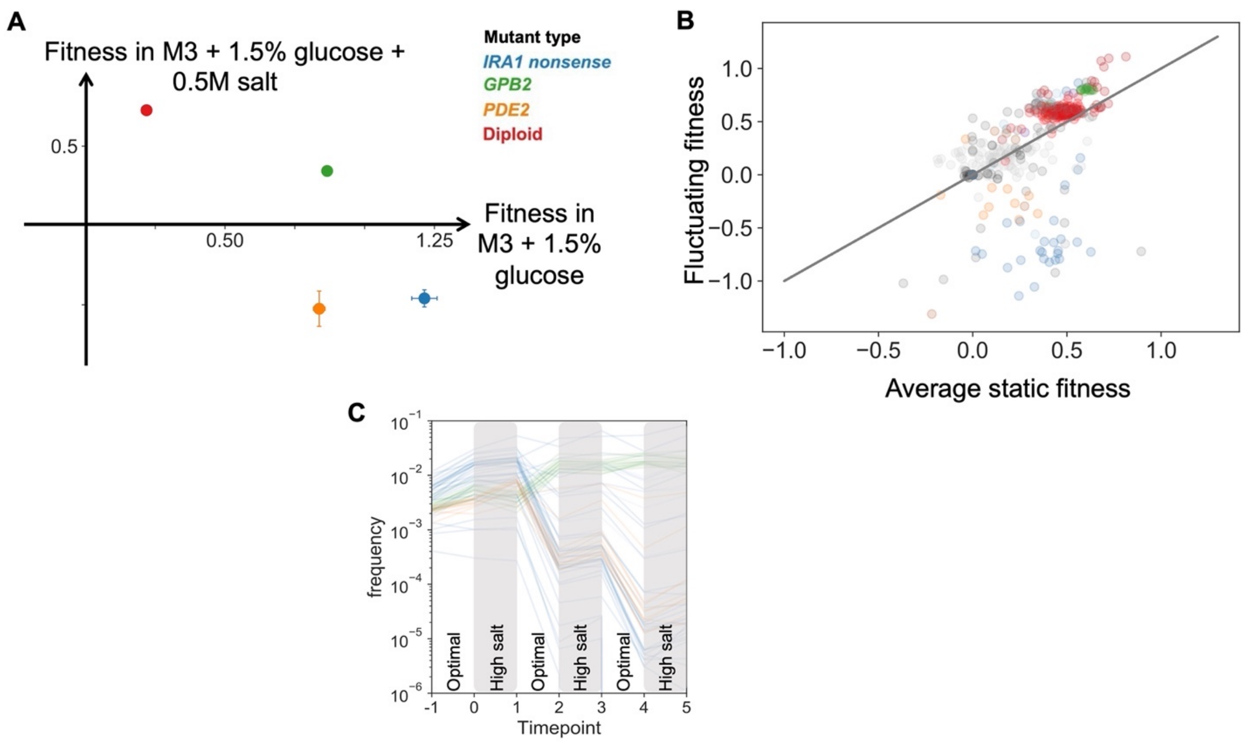


**Figure S9: Environmental memory observed in IRA1, PDE2 mutants in high-salt fluctuations.** **A** We re-measured fitness of mutants from a previous study[12], finding that they have antagonistic tradeoffs in two static environments. **B** We also measured fitness in a fluctuating environment alternating between these two conditions. We find that the mutants with antagonistic tradeoffs, those with SNPs in the IRA1 and PDE2 genes, exhibit non-additivity. **C** Examining the frequency trajectories of these mutants (orange and blue) reveals that they crash during the optimal cycle in which they should be highly adaptive, and stagnate during the cycle in which they should be deleterious. This result is a clear example of environmental memory, since the fitness in each component differs dramatically from the fitness in the corresponding static environment.

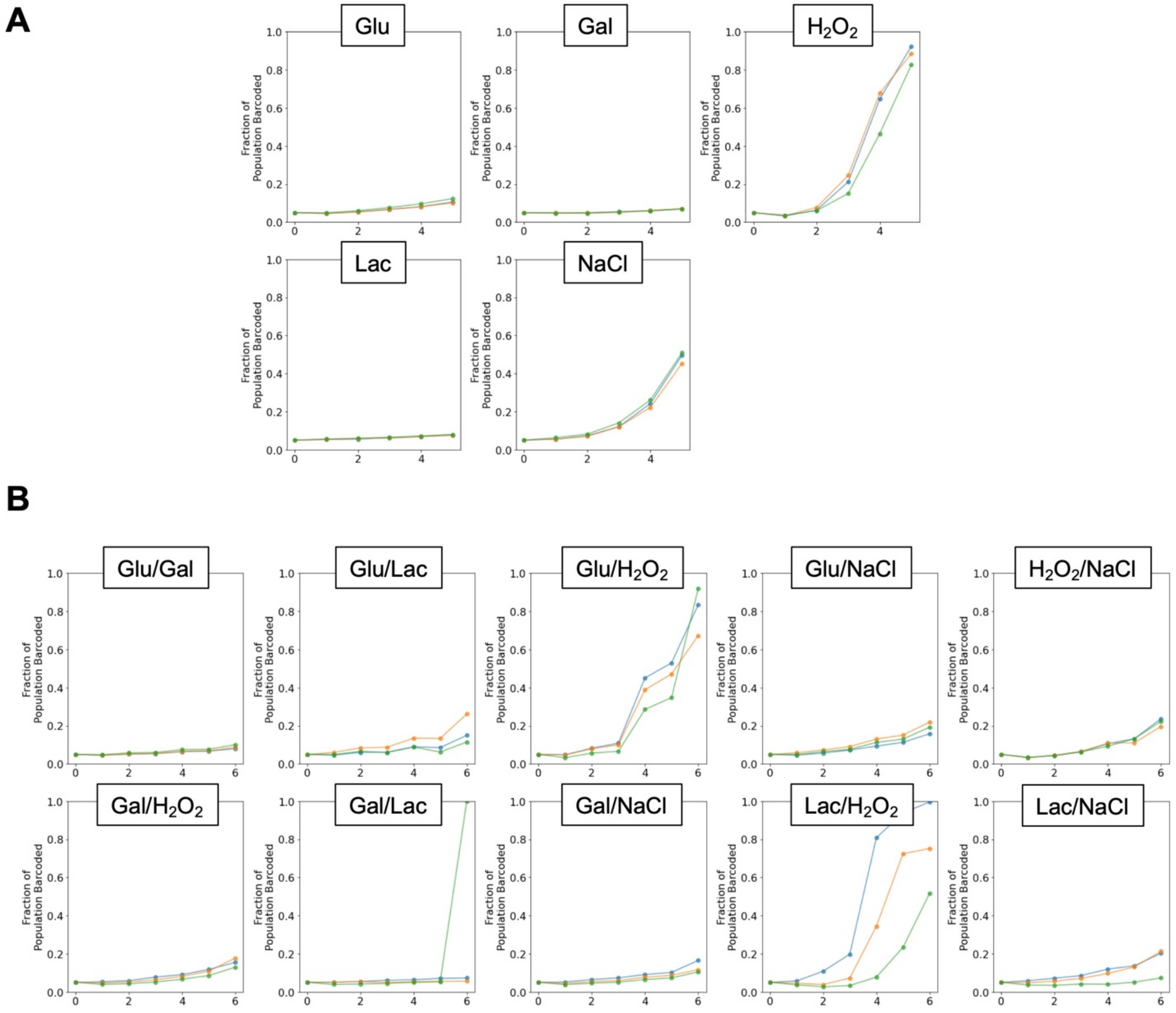


**Figure S10: Change in fraction of population occupied by barcoded mutant pool over time in each fitness measurement.** We calculated the fraction of the population occupied by the barcoded pool in the static (panel **A**) and fluctuating (panel **B**) environment fitness measurements as in ref [3]. In order to avoid frequency-dependent effects and measure fitness while in competition with the ancestor rather than among the pool, we discarded timepoints in which all three replicates (shown in different colors) reached frequencies greater than 50%, including the final timepoints of *$H_2O_2$*, *Glu*/*$H_2O_2$*, and *Lac*/*$H_2O_2$*. (The final timepoint in one of the *Gal*/*Lac* replicates was not successfully sequenced, and sequencing error attributes a single barcode to this timepoint, which we did not use.)

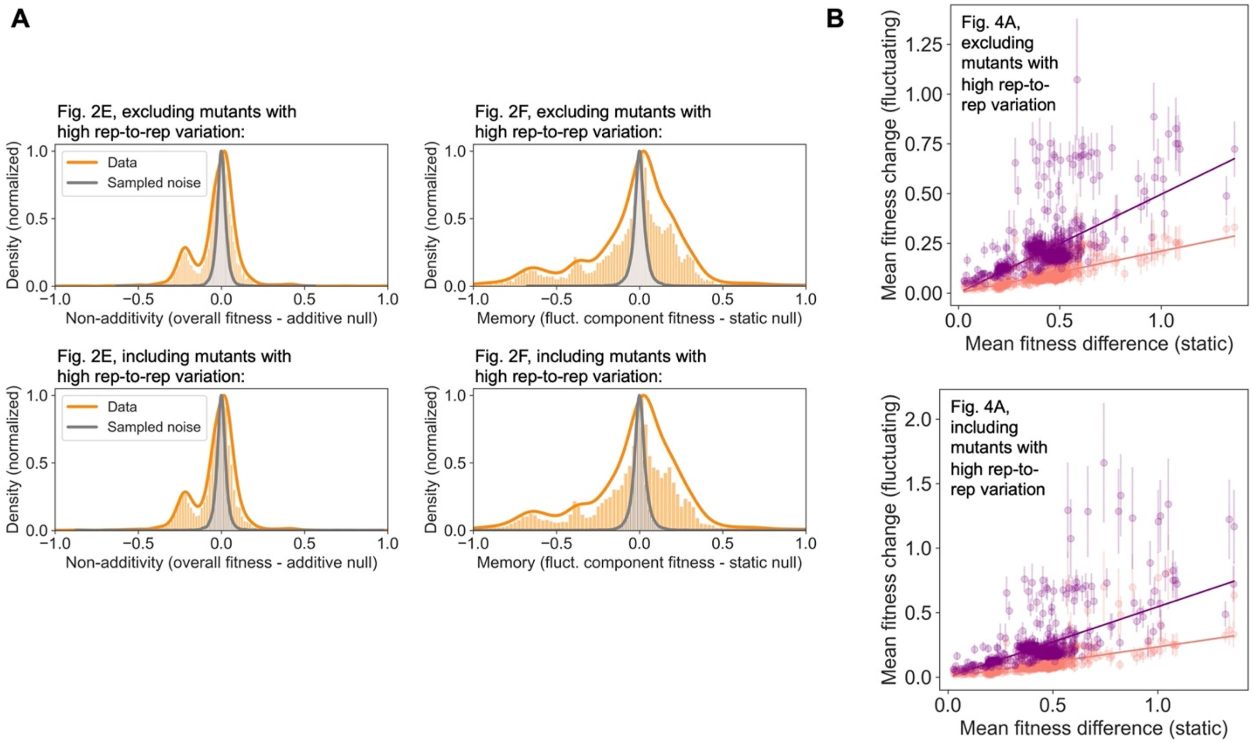


**Figure S11: Trends do not change when including mutants with high replicate-to-replicate variation.** We discarded 12 mutants from our analyses that have average replicate-to-replicate difference > 0.2, but including them does not change the final results from Fig. 2 (panel **A**) and Fig. 4 (panel **B**).

**Supplementary Discussion**

**Supplementary Note 1: Mathematical model shows that differing yield alone cannot explain observed non-additivity and memory**

In the main text, we observe that both non-additivity and memory correlate with fitness difference (Fig. 4A, main text). We define non-additivity as the difference between fitness in a fluctuating environment and the average of fitness in the static environment components. We define memory as the difference between fitness in a single component of a fluctuating environment and fitness in the corresponding static environment.

Our fluctuating environments are composed of pairwise combinations of five static environments. In one of the five static environments (*Lac*, which contains lactate as the carbon source), we observed that population yield is about half of the yield reached in all other environments. Below, we demonstrate with a mathematical model that differences in yield alone are not sufficient to explain the non-additivity and memory that we observe in our data.

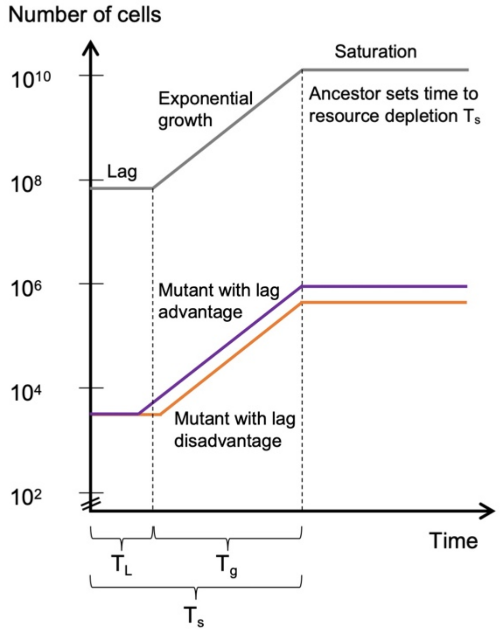


**Figure 1**: In the model, the ancestor (gray) sets the time to resource depletion, before which mutants can grow.

Model description

This model is similar to a consumer resource model, but simplified to three static phases—lag, exponential phase, and stationary phase—which are assumed to be no growth, exponential growth, and no growth (**Fig. 1**, right).

We assume the ancestor dominates the population and that mutants occupy a negligible fraction, so that the time to stationary phase, $T_s$, is set by the time it takes the ancestor to consume the carbon source. After the cycle is over, the population is diluted by a dilution factor DF and the phases begin again.

The fitness $s_i$ of mutant $i$ is defined as:

$$s_i = \ln\left(\frac{n_{i1}}{n_{i0}}\right) - \ln\left(\frac{n_{a1}}{n_{a0}}\right) \tag{1}$$

where $n_{i1}$ is the number of cells of the mutant $i$ after one cycle and $n_{i0}$ is the number of cells at the start of the cycle, and similarly for $n_a$, the number of ancestor cells. The ratio of ancestor cells will always be the same if the dilution factor is the same, because it dominates the population:

$$\ln\left(\frac{n_{a1}}{n_{a0}}\right) = \ln(DF) \tag{2}$$

In our experiments, the dilution factor DF is 250 (representing about eight generations of growth). Given that the ancestor can reach saturation during the cycle, this ratio will always be equal to the dilution factor, regardless of the saturation yield—a low-yield environment will lead to a smaller bottleneck population size, but the same ratio of final and initial numbers.

<u>Growth of ancestor in fluctuating conditions</u>
In a fluctuating environment, however, a fluctuating yield will cause the ratio to fluctuate, because when the lower-yield environment is diluted into the higher-yield environment, the smaller bottleneck population will grow more than eight generations, and vice versa. Assuming environment A is followed by a dilution of factor DF into environment B, we can calculate the number of doublings $d$ in environment B in terms of the yields $y$ of environments A and B:

$$2^{d_B} = \frac{y_B}{y_A} DF \tag{3}$$

If the yields are equal, the doublings will be determined by the dilution factor alone. If not, we must account for the number of doublings in each environment:

$$d_B = \frac{\ln\left(\frac{y_B DF}{y_A}\right)}{\ln(2)} \qquad d_A = \frac{\ln\left(\frac{y_A DF}{y_B}\right)}{\ln(2)} \tag{4}$$

The ratio of ancestor cells in environment A then becomes:

$$\ln\left(\frac{n_{a1}}{n_{a0}}\right) = \ln(2^{d_A}) = \ln\left(2^{\frac{\ln\left(\frac{y_A DF}{y_B}\right)}{\ln(2)}}\right) = \ln\left(exp\left(ln\left(\frac{y_A DF}{y_B}\right)\right)\right) = \ln\left(\frac{y_A}{y_B} DF\right) \tag{5}$$

If the yields are equal, Eqn. 5 reduces to Eqn. 2.

<u>Fitness of a mutant in a static environment</u>
This is true for the ancestor, but what about a mutant $i$? In a static environment, the ancestor takes some time $T_s$ to reach saturation. This time is made up of periods of lag and growth:

$$T_s = T_L + T_g \tag{6}$$

$T_g$ can be expressed in terms of the ancestor's growth rate $r_a$:

$$T_g = \frac{1}{r_a} \ln(DF) \tag{7}$$

The mutant also grows during time $T_g$, and can potentially grow during the ancestor's lag time; alternatively, the ancestor may grow during the mutant's lag time, depending on who has a longer lag. So we write the change in the number of mutant cells in terms of both and plug in Eqn. 7:

$$\frac{n_{i1}}{n_{i0}} = \exp\left(r_i\left(T_g + (T_L - T_{Li})\right)\right) = \exp\left(r_i(\frac{1}{r_a}\ln(DF) + \Delta T_L\right) = \exp(r_i\Delta T_L)\left(DF^{\frac{r_i}{r_a}}\right) \quad (8)$$

where $\Delta T_L = T_L - T_{Li}$, the mutant's lag (dis)advantage.

Now we can plug Eqns. 2 and 8 into Eqn. 1:

$$s_i = \ln\left(\frac{n_{i1}}{n_{i0}}\right) - \ln\left(\frac{n_{a1}}{n_{a0}}\right) = \ln\left(exp(r_i\Delta T_L)\left(DF^{\frac{r_i}{r_a}}\right)\right) - \ln(DF) = r_i\Delta T_L + \left(\frac{r_i - r_a}{r_a}\right)\ln(\mathrm{DF}) \quad (9)$$

This divides the mutant's fitness into its lag and growth (dis)advantages over the ancestor. If both are equal to the ancestor, the fitness is zero. We can try plugging in some realistic numbers: dilution factor 250, ancestor growth rate 0.4 (doubling time of ~90 min), and an adaptive mutant with a growth rate of 0.45 and a lag time advantage of two hours:

$$s_i = 0.45 * 2 + \left(\frac{0.45 - 0.4}{0.4}\right)\ln(250) = 1.59 \quad (10)$$

This in the range of expected per-cycle fitness of a highly adaptive mutant. Notice that a two-hour lag advantage, which seems reasonable, contributes more to fitness gain than a 12.5% increase in growth rate, which seems high. Decreasing lag seems like an easier way to gain fitness than increasing growth rate, and could even still pay off with a decreased growth rate.

<u>Fitness of a mutant in a fluctuating environment</u>

The above calculation applies to any static environment where the ancestor can reach saturation, regardless of yield. But in a fluctuating environment, will differing yields lead to non-additive fitness?

We can modify Eqn. 9 for a fluctuating environment with differing yields as we did in Eqn. 5, by multiplying instances of $DF$ by $\frac{y_A}{y_B}$ for environment A, or $\frac{y_B}{y_A}$ for environment B:

$$s_{iA} = r_{iA}\Delta T_{LA_f} + \left(\frac{r_{iA} - r_{aA}}{r_{aA}}\right)\ln\left(\frac{\mathrm{y_A}}{\mathrm{y_B}}\mathrm{DF}\right) \quad (11)$$

And similarly for environment B:

$$s_{iB} = r_{iB}\Delta T_{LB_f} + \left(\frac{r_{iB} - r_{aB}}{r_{aB}}\right)\ln\left(\frac{\mathrm{y_B}}{\mathrm{y_A}}\mathrm{DF}\right) \quad (12)$$

The average fluctuating fitness $\bar{s}_f$ is then:

$$\bar{s}_f = \frac{\left(r_{iA}\Delta T_{LA_f} + \left(\frac{r_{iA} - r_{aA}}{r_{aA}}\right)\ln\left(\frac{\mathrm{y_A}}{\mathrm{y_B}}\mathrm{DF}\right) + r_{iB}\Delta T_{LB_f} + \left(\frac{r_{iB} - r_{aB}}{r_{aB}}\right)\ln\left(\frac{\mathrm{y_B}}{\mathrm{y_A}}\mathrm{DF}\right)\right)}{2} \quad (13)$$

To find the non-additivity, we subtract the mean fitness across the static environments from the mean fluctuating fitness:

$$\bar{s}_f - \bar{s}_s = \frac{\left(r_{iA}\Delta T_{LA_f} + \left(\frac{r_{iA} - r_{aA}}{r_{aA}}\right)\ln\left(\frac{\mathrm{y_A}}{\mathrm{y_B}}\mathrm{DF}\right) + r_{iB}\Delta T_{LB_f} + \left(\frac{r_{iB} - r_{aB}}{r_{aB}}\right)\ln\left(\frac{\mathrm{y_B}}{\mathrm{y_A}}\mathrm{DF}\right)\right)}{2} - \frac{r_{iA}\Delta T_{LA_s} + \left(\frac{r_{iA} - r_{aA}}{r_{aA}}\right)\ln(\mathrm{DF}) + r_{iB}\Delta T_{LB_s} + \left(\frac{r_{iB} - r_{aB}}{r_{aB}}\right)\ln(DF)}{2}$$

$$= \frac{r_{iA}\left(\Delta T_{LA_f} - \Delta T_{LA_s}\right) + r_{iB}\left(\Delta T_{LB_f} - \Delta T_{LB_s}\right) + \left(\frac{r_{iA} - r_{aA}}{r_{aA}}\right)\left(\ln\left(\frac{\mathrm{y_A}}{\mathrm{y_B}}\mathrm{DF}\right) - \ln(\mathrm{DF})\right) + \left(\frac{r_{iB} - r_{aB}}{r_{aB}}\right)\left(\ln\left(\frac{\mathrm{y_B}}{\mathrm{y_A}}\mathrm{DF}\right) - \ln(\mathrm{DF})\right)}{2}$$

$$= \frac{r_{iA}\left(\Delta T_{LA_f} - \Delta T_{LA_s}\right) + r_{iB}\left(\Delta T_{LB_f} - \Delta T_{LB_s}\right) + \ln\left(\frac{y_A}{y_B}\right)\left(\left(\frac{r_{iA} - r_{aA}}{r_{aA}}\right) - \left(\frac{r_{iB} - r_{aB}}{r_{aB}}\right)\right)}{2}$$

$$= \frac{r_{iA}\Delta T_{LA} + r_{iB}\Delta T_{LB} + \ln\left(\frac{y_A}{y_B}\right)\delta r}{2} \tag{14}$$

In the last line of Eqn. (14), we have abbreviated some terms to delineate the two mechanisms that can lead to non-additivity: changing yields and changing lags.

First, $\delta r = \left(\frac{r_{iA} - r_{aA}}{r_{aA}}\right) - \left(\frac{r_{iB} - r_{aB}}{r_{aB}}\right)$ is the difference in growth advantage across the two static environments. This term only applies if the ancestor's yield in the two environments is different, and in that case, we expect to see non-additivity arising from this difference.

Second, $\Delta T_{LA} = \Delta T_{LA_f} - \Delta T_{LA_s}$ and $\Delta T_{LB} = \Delta T_{LB_f} - \Delta T_{LB_s}$ represent changes in lag (dis)advantages of the mutant compared to the ancestor in fluctuating conditions. If $\Delta T_{LA} > 0$, the mutant's lag time in environment A is more advantageous, as compared to the ancestor, when it is fluctuating from environment B than when it is growing in static environment A. Since the lag advantage is defined in comparison to the ancestor, $\Delta T_{LA} > 0$ could mean that the ancestor increases its lag in fluctuating conditions more than the mutant, or that mutant decreases its lag more than the ancestor, or any other reason causing the mutant's lag compared to the ancestor to improve in fluctuating conditions. These and any other changes to the lag compared to the ancestor in either environment A or B in fluctuating vs. static conditions will lead to non-additivity.

<u>Unequal yields alone cannot explain non-additivity in data</u>

As a first step toward explaining non-additivity in our data, we assume that in a fluctuating environment, the lag time of the mutant changes in the same way as the lag time of the ancestor. This means that changes in lag time will not cause non-additivity, because $\Delta T_{LA_f} = \Delta T_{LA_s}$ and

$\Delta T_{LB_f} = \Delta T_{LB_s}$, and $\Delta T_{LA} = \Delta T_{LB} = 0$. Under this assumption, the only mechanism that could cause non-additivity is differing yield. Eqn. (14), with $\Delta T_{LA} = \Delta T_{LB} = 0$, determines the non-additivity:

$$\bar{s}_{f|\Delta T_L=0} - \bar{s}_{s|\Delta T_L=0} = \frac{\ln\left(\frac{y_A}{y_B}\right)\left(\left(\frac{r_{iA} - r_{aA}}{r_{aA}}\right) - \left(\frac{r_{iB} - r_{aB}}{r_{aB}}\right)\right)}{2} \qquad (15)$$

The non-additivity scales with the difference in fitness across static environments, which is proportional to the expression in the parentheses of the numerator. This means that as difference in fitness grows, so does non-additivity—the same direction of the effect we observe in our data (Fig. 4A, main text).

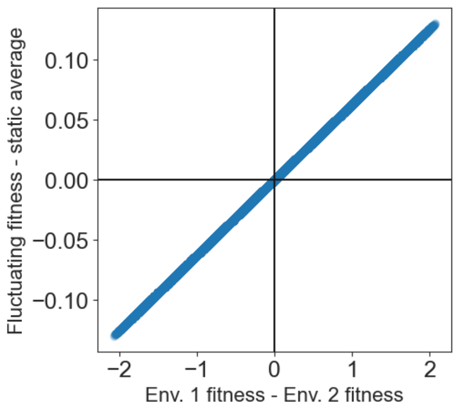


**Figure 2**: Non-additivity caused only by changing yields in a fluctuating environment.

If we plot Eqn. (15) with $y_A/y_B = 2$, $r_{aA} = 0.4$, $r_{aB} = 0.2$, and $r_{iA}$ and $r_{iB}$ spanning a range centered around these quantities, we get the result plotted in **Fig. 2** to the right. The x-axis is difference in fitness across static environments, or $\log(DF)$ multiplied by the expression in the parentheses in the numerator of Eqn. (15). The y-axis is the non-additivity, and it's clear that a two-fold difference in yield across fluctuating environments can generate significant non-additive effects for these parameters. The sign of the non-additivity is determined by the sign of fitness difference (unless $\ln\left(\frac{y_A}{y_B}\right) < 0$, which reverses the sign), so the trend will be monotonic (increasing if $\ln\left(\frac{y_A}{y_B}\right) > 0$ or decreasing if $\ln\left(\frac{y_A}{y_B}\right) < 0$). The trend passes through the origin, and the trend has a one-to-one mapping of fitness difference to

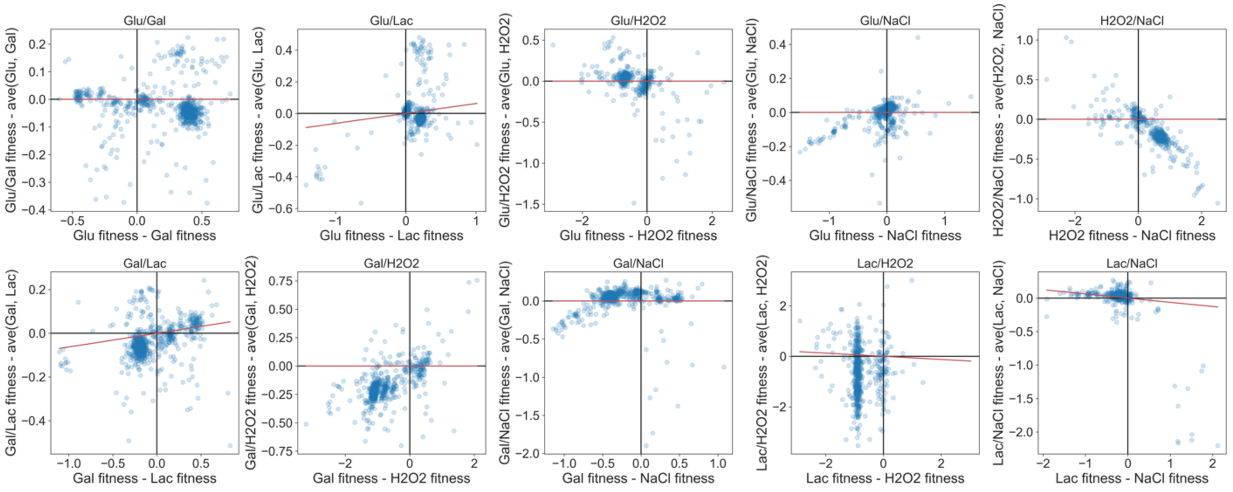


**Figure 3**: Data from ten fluctuating environments does not fit model the of non-additivity due to changing yield alone (red lines). While some data does appear to fit the trend, there are highly non-additive data points sitting far from the line (note that the scale in the *Lac/NaCl* plot makes data look closer to the line).

non-additivity. These constraints provide guidelines for interpreting real data: in order to be explained by differing yields alone, a trend must 1) be monotonic; 2) pass through the origin; 3) have a 1-1 mapping (only error can add width to the line).

Let's look at real data. In **Fig. 3**, we plot fitness difference vs. non-additivity, using real rather than absolute values. The only environment in which we measured a difference of yield was the lactate (*Lac*) environment, which grew to about half the number of cells as the rest of the environments (based on OD readings during evolution experiments). Using this difference, we have plotted the expected trendlines (red) in all pairs containing *Lac*. In the other plots, we would expect trendlines with a slope of zero. In two out of these four pairs (*Gal*/*Lac*, *Lac*/*NaCl*), some of the data does fall roughly near the trendline, suggesting that the effect of variable yield can be seen. However, many data points still sit far from the line (and note the large scale in the *Lac*/*NaCl* plot). This data appears to rule out the hypothesis that differing yields alone can explain our observed non-additivity.

<u>Unequal yields alone cannot explain memory in data</u>

To see the effect that unequal yields have on fitness changes in individual components of fluctuating environments, we can compare the fitness in an environment when it is part of a fluctuating environment with differing yields to when it is part of a static environment with equal yields. Subtracting Eqn. 9 from Eqn. 11 gives the expected change in fitness, or memory, due to this effect:

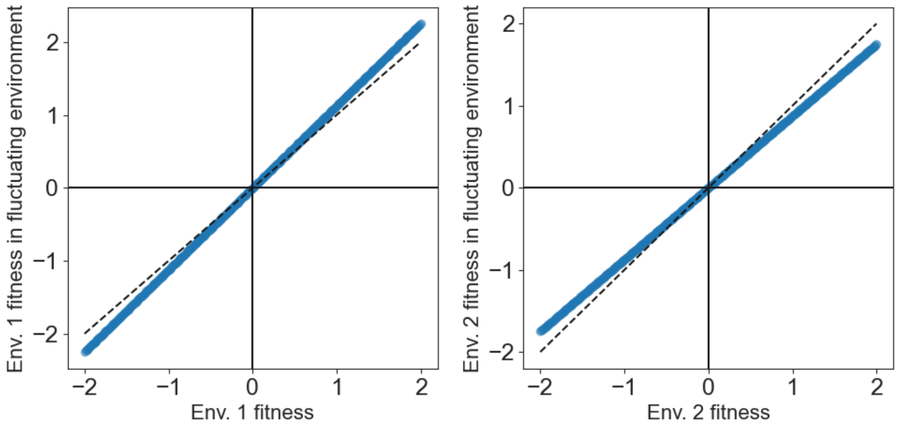


**Figure 4**: Unequal yields lead to memory in individual components of a fluctuating environment. The memory is the deviation of the simulated data (blue) from the dashed line.

$$s_{iAf} - s_{iAs} = \left(\frac{r_{iA} - r_{aA}}{r_{aA}}\right)\left(\ln\left(\frac{\mathrm{y_A}}{\mathrm{y_B}}\mathrm{DF}\right) - \ln(DF)\right)$$

$$= \left(\frac{r_{iA} - r_{aA}}{r_{aA}}\right)\ln\left(\frac{\mathrm{y_A}}{\mathrm{y_B}}\right) \quad (16)$$

This equation can be re-written in terms of $s_{iAs}$, the fitness in the static environment A:

$$s_{iAf} = s_{iAs}\left(1 + \frac{\ln\left(\frac{y_A}{y_B}\right)}{\ln(DF)}\right) \quad (17)$$

When the yields in environments A and B are equal, the term in parentheses goes to 1, and the fluctuating component fitness is equal to the fitness in the static environment, and there is no memory. In **Fig. 4** above, we plot Eqn. 17 with $DF = 250$ and $y_A/y_B = 2$. The data is shown in blue, and the dashed line represents the equal-yield null model with no memory. Memory can thus be seen from deviations from this dashed line.

In **Fig. 5** to the right, we show real data. These data reveal two interesting patterns. First, fitness often departs dramatically from the dashed line, indicating that memory is widespread and strong.
Second, in fluctuating environments with unequal yields (plots with red lines), the expected memory (red line) is not able to explain these deviations. Even in a fluctuating environment with a two-fold difference in yield across environments, the expected memory is subtle compared to what we observe in our actual data.

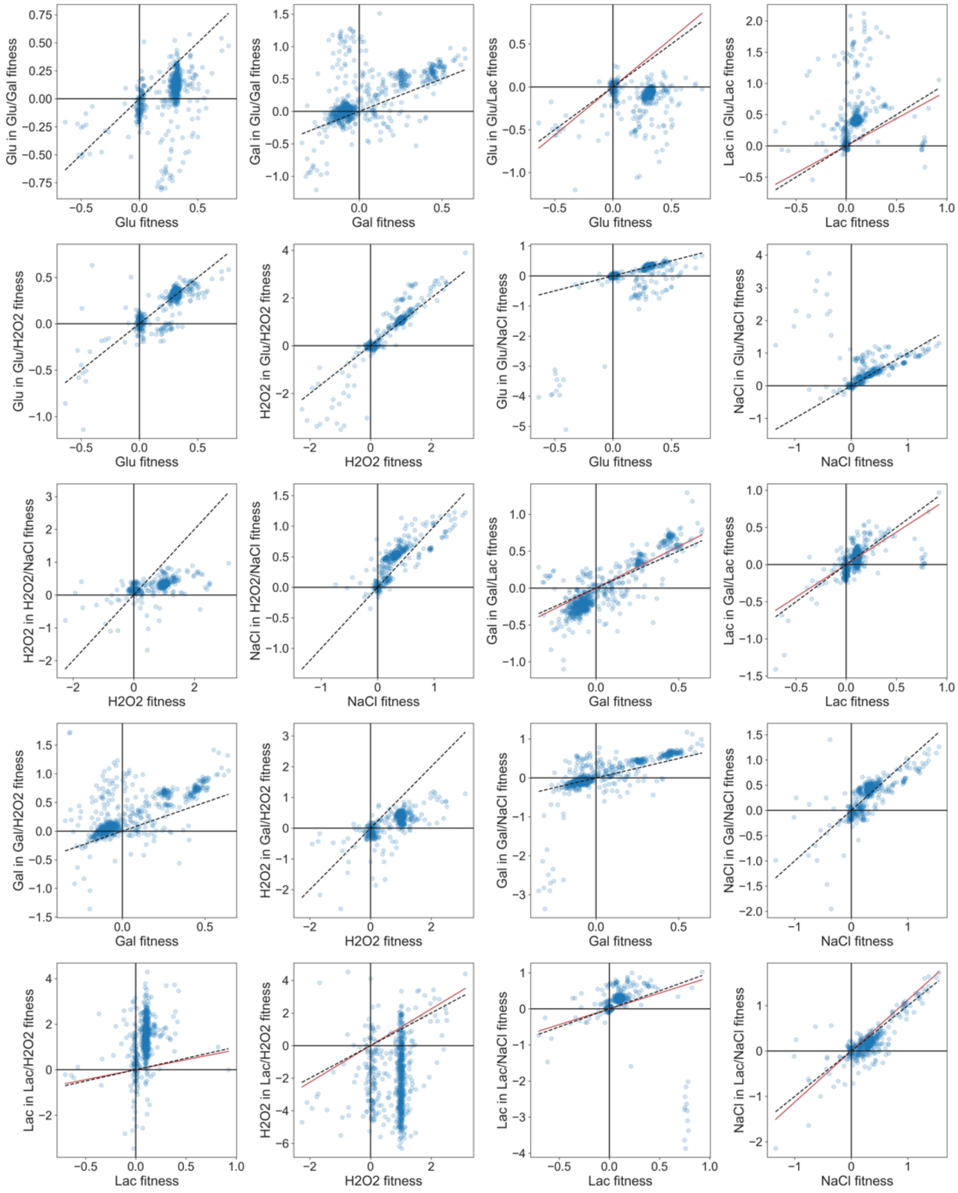


**Figure 5**: Memory in our data appears as deviations from the dashed lines. Unequal yields would cause slight deviations (red lines) that cannot explain the amount of memory that we observe.

Altogether, Fig. 3 and Fig. 5 demonstrate that unequal yield in fluctuating environments that contain the *Lac* condition cannot explain the non-additivity and memory that we observe in our data. Changes in lags and/or growth rates in fluctuating environments are most likely contributing to these effects.
